# The protein map of the protozoan parasite *Leishmania (Leishmania) amazonensis*, *Leishmania* (*Viannia) braziliensis* and *Leishmania (Leishmania) infantum* during growth phase transition and temperature stress

**DOI:** 10.1101/2023.11.27.568882

**Authors:** Simon Ngao Mule, Joyce Silva Saad, Ismael Pretto Sauter, Livia Rosa Fernandes, Gilberto Santos de Oliveira, Daniel Quina, Fabia Tomie Tano, Deborah Brandt-Almeida, Gabriel Padrón, Beatriz Simonsen Stolf, Martin R. Larsen, Mauro Cortez, Giuseppe Palmisano

**Author notes:** These authors contributed equally (Simon Ngao Mule, Joyce Silva Saad). To whom correspondence should be addressed Prof. Giuseppe Palmisano Prof. Mauro Cortez.

## Abstract

*Leishmania* parasites cause a spectrum of diseases termed leishmaniasis, which manifests in two main clinical forms, cutaneous and visceral leishmaniasis. *Leishmania* promastigotes transit from proliferative exponential to quiescent stationary phases inside the insect vector, a relevant step that recapitulates early molecular events of metacyclogenesis. During the insect blood meal of the mammalian hosts, the released parasites interact initially with the skin, an event marked by temperature changes. Deep knowledge on the molecular events activated during *Leishmania*-host interactions in each step is crucial to develop better therapies and to understand the pathogenesis. In this study, the proteomes of *Leishmania* (*Leishmania) amazonensis* (La), *Leishmania* (*Viannia) braziliensis* (Lb), and *Leishmania* (*Leishmania) infantum* (syn *L. L. chagasi)* (Lc) were analyzed using quantitative proteomics to uncover the proteome modulation in three different conditions related to growth phases and temperature shifts: 1) exponential phase (Exp); 2) stationary phase (Sta25) and; 3) stationary phase subjected to heat stress (Sta34). Functional validations were performed using orthogonal techniques, focusing on α-tubulin, gp63 and heat shock proteins (HSPs). Species-specific and condition-specific modulation highlights the plasticity of the *Leishmania* proteome, showing that pathways related to metabolism and cytoskeleton are significantly modulated from exponential to stationary growth phases, while protein folding, unfolded protein binding, signaling and microtubule-based movement were differentially altered during temperature shifts. This study provides an in-depth proteome analysis of three *Leishmania* spp., and contributes compelling evidence of the molecular alterations of these parasites in conditions mimicking the interaction of the parasites with the insect vector and vertebrate hosts.

## INTRODUCTION

Leishmaniasis is a group of neglected vector-borne diseases caused by the flagellate protozoan *Leishmania*, which manifests in different clinical forms according to the infecting species and the host’s immune response [1, 2]. It may present as asymptomatic or in the two primary forms: cutaneous leishmaniasis (CL) and visceral leishmaniasis (VL) [3]. Cutaneous leishmaniasis may present as localized, which is the most common form of CL, and is associated with *Leishmania braziliensis* and *Leishmania amazonensis* infections in Brazil, or the rare diffuse-CL caused by *L. amazonensis*, and the disseminated CL and mucocutaneous leishmaniasis, associated with *Leishmania braziliensis* [4-6]. Complications during diffuse leishmaniasis have shown that this form caused by *Leishmania amazonensis* has a more severe course and does not respond well to antimonial treatment [7], whereas mucocutaneous leishmaniasis by *Leishmania braziliensis* is associated with a progressive mutilating disease that is difficult to heal [3]. Visceral leishmaniasis, also called kala-azar, is more aggressive than cutaneous leishmaniasis, since the infection affects several organs or tissues such as the spleen, liver, and bone marrow. In Brazil, the main species responsible for cutaneous leishmaniasis is *L.* (*Viannia*) *braziliensis*, followed by *L.* (*Leishmania*) *amazonensis* [3], while visceral leishmaniasis is caused by *L.* (*Leishmania*) *chagasi* [8].

*Leishmania* biological cycle goes through three main different parasite life stages: procyclic promastigote, found in the exponential proliferative phase during parasite development within the insect vector (phlebotomine sand fly); metacyclic promastigote that is transmitted to the mammalian host during a blood meal of female phlebotomine sand fly, and; amastigote, an intracellular form derived from the transition of metacyclic promastigotes within phagocytic cells, such as macrophages [9]. The molecular alterations during the biological cycle transition constitute an important opportunity to understand the underlined *Leishmania* biology and for diagnostic or therapeutic applications [10]. Parasite factors associated with infectivity, virulence and establishment of the disease can be found differentially expressed according to the developmental stage, helping the parasite to survive within the insect and vertebrate hosts [11, 12]. Among these factors, lipophosphoglycan (LPG) and gp63 are examples of such molecules found in the surface membrane of the parasite. LPG is a species-specific glycoconjugate [13, 14], which undergoes structural modifications during metacyclogenesis, such as elongation of the phosphorylated glycan units and change in the terminal glycans from galactose to arabinopyrose [15, 16]. These changes assist in the metacyclic promastigote opsonization via the complement system, favoring macrophage invasion and avoiding parasite lysis [17, 18], contrary to procyclic promastigotes which are rapidly lysed in the presence of serum [19, 20]. Moreover, LPG levels are dramatically reduced in amastigotes [21]. The stage-specificity of LPG has been demonstrated in other species that cause cutaneous leishmaniasis, such as *L. major*, where amastigotes have modifications with relatively larger phosphoglycan units than promastigotes LPG [22, 23]. Gp63, also known as zinc-metalloprotease 63 or leishmanolysin, is a surface antigen highly expressed in promastigote and in lower levels in amastigote stages [24] and protects the parasite against earlier infection phases in the mammalian host [25] by inhibiting the innate immune response. Furthermore, this metalloprotease acts by preventing parasite lysis by the complement system [26], assists parasite adherence to macrophages [27], and degrades the extracellular matrix favoring its migration [25].

In order to survive and proliferate during its digenetic biological cycle, *Leishmania* parasites remodel their proteome in response to external stimuli imposed by the host, where the temperature plays a key role on the different parasite environment. More important, each species has a different and optimal temperature for its development [28]. *Leishmania* species associated with VL or CL from the Old World do not show significant differences in their growth, developing well in temperatures between 30 to 32 °C; however, for species from the New World associated with VL like *L. chagasi*, the parasites proliferate better in temperatures above 32°C than the CL species. Moreover, the most resistant to higher temperatures between CL species are *L. mexicana*, followed by *L. braziliensis* and lastly *L. amazonensis* [29]. Despite this difference, *L. mexicana* and *L. amazonensis* are classified within the same subgenus and complex (*mexican* complex), suggesting that temperature sensitivity is not correlated to the clinical manifestation caused by the New World species’ [29]. Among the molecular factors involved in resistance to temperature stress, the heat shock proteins (HSPs) play a key role in parasite differentiation and survival, increasing its abundance during differentiation of promastigotes to amastigotes [30-32].

*Leishmania* parasites regulate their gene expression at post-transcriptional and post-translational levels, making it difficult to find a relationship between their transcriptome and proteome [12, 33]. Proteomics has been used to create *Leishmania* protein maps to corroborate and complement genomic sequencing studies [33]. For example, one of the first proteomic studies in *Leishmania* compared promastigotes and axenic amastigotes of *L. infantum*, observing 62 differentially expressed proteins [34]. Another study using *L. panamensis* reported 45 and 75 proteins differentially expressed in promastigotes and axenic amastigotes, respectively [35]. Recently, studies have reported using MALDI-TOF MS to identify *Leishmania* species from *in vitro* cultures [36, 37]. In addition, the application of mass-spectrometry-based proteomics to leishmaniasis has been recently reviewed, reinforcing its application for disease therapeutics, biomarker discovery, drug resistance, and identification of vaccine candidates [10, 38, 39].

In this study, we performed a quantitative mass spectrometry-based proteomics to compare the proteome of three *Leishmania* species from the New World: *L. amazonensis* (La)*, L. braziliensis* (Lb) and *L. infantum (syn L. chagasi)* (Lc) in three conditions: 1) exponential phase at 25°C (Exp); 2) stationary phase at 25°C (Sta25) and; 3) stationary phase subjected to heat stress at 34°C (Sta34). In particular, the first stationary phase (Sta25) was cultured and maintained at 25°C for 7 days, while the Sta34 was maintained at 25°C for 6 days and then shifted to 34°C for 24 hours. This last condition mimics the thermal shock when the parasites are transferred from the phlebotomine sand fly to the vertebrate host. Differentially expressed HSPs, cytoskeleton, and GP63 proteins were assayed by western blotting, zymography, and RT-qPCR. Moreover, temperature sensitivity assays were performed for the three *Leishmania* species. This study interrogates for the first time the proteome of three *Leishmania* species from the New World and its modulation in different conditions ranging from two different growth stages and temperatures. The results reveal important insights into the species-specific molecular changes of *Leishmania* parasites exposed to different stimuli.

## MATERIALS AND METHODS

### Parasite cell culture

*L. amazonensis* (IFLA/BR/67/PH8*), L. braziliensis* (MHOM/BR/75/M2903), and *L. infantum (syn L. chagasi)* (MHOM/BR/1972/LD) promastigotes were cultured as described previously [40, 41], with some modifications. Briefly, parasites were incubated in M199 medium (Vitrocell) supplemented with 20% inactivated fetal bovine serum (FBS) (Vitrocell), 40 mM HEPES, 2.5 µg/mL hemin, 10 mM adenine, 2 mM L-glutamine, 2 µg/mL D-biotin, 100 U/mL penicillin, 100 µg/mL streptomycin, 2% filter sterilized human urine, at pH 7.4. Proteomes from three different conditions were analysed in this study. For the first analysis, the exponential/logarithmic and stationary growth phases were collected on the 3^rd^ and 7^th^ days of culture at 25°C, respectively. For the second analysis, stationary phase promastigotes on the 6^th^ day of culture at 25°C were grown (or not) at 34°C for 24 hours and collected on the 7^th^ day (**Fig. 1**). The parasites were collected separately by centrifugation at 2100 g for 10 min at 4°C, washed twice with phosphate-buffered saline (PBS: 137 mM NaCl, 2.7 mM KCl, 10 mM Na_2_HPO_4_, 1.8 mM of KH_2_PO_4_, pH 7.4), and pelleted by centrifugation as previously described. All analyses were done in three biological replicates.

**Fig. 1.**
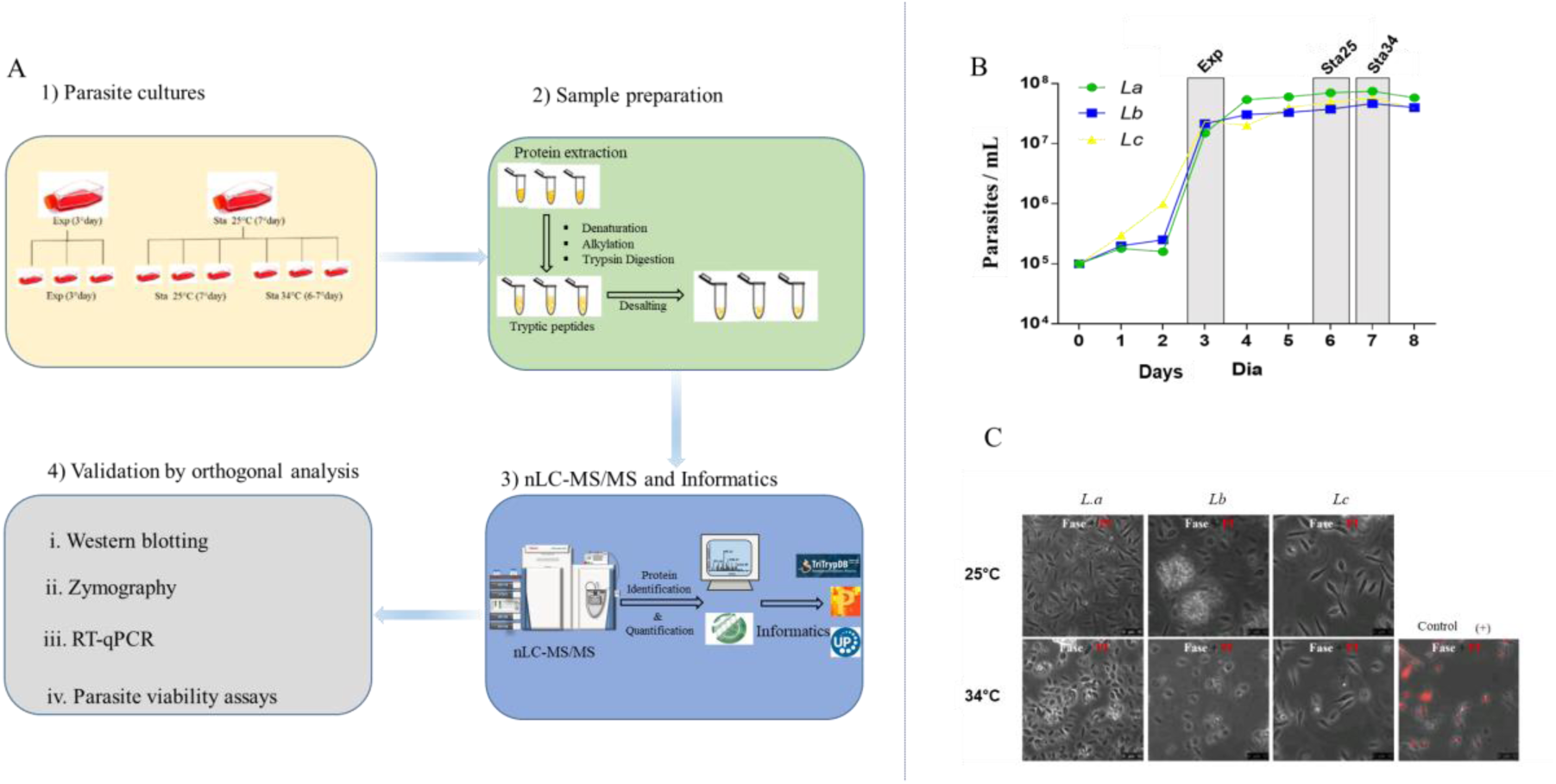
Experimental workflow of proliferative and viable parasites to study *Leishmania amazonensis*, *Leishmania braziliensis* and *Leishmania infantum* proteome modulation in different conditions. **A**) *Leishmania* parasites were collected at different growth phases and temperature shift: exponential (Exp), stationary 1 (Sta25), and stationary 2 (Sta34) as described in the Materials and Methods. Proteins were extracted, digested with trypsin, and analysed using nanoLC-MS/MS for protein identification and quantification. Selected protein candidates were validated using orthogonal techniques such as western-blot, MTT, RT-qPCR, and Zymography. **B**) Growth curve of the promastigote form of the three species over eight days. Parasite cells in the exponential phase were collected on day 3 (Exp), and stationary phases Sta25 and Sta34 on day 7. The stationary phase, Sta34, was submitted to a temperature of 34°C for 24 hours while Sta25 was maintained at 25°C. The curve is a representative result of two independent assays. La = *L. amazonensis* (green), Lb = *L. braziliensis* (blue) and Lc = *L. infantum chagasi* (yellow). C) Viability assay with propidium iodide. The parasites were treated with 0.4% PFA and 10 μg/mL PI for 5 min. DMSO-treated parasites (20% v/v) was used as (+) control for cell death.

### Morphometry of parasite forms from stationary phase

To validate the proper synchronization of the different forms in each parasite culture, we analyzed and quantified the number of metacyclic promastigotes during the stationary phase, which define the phenotypic similarities between the different species culture. The parasites were stained with panoptic stain, which allow us to visualize different structures such as body, flagellum, nucleus and kinetoplast of the parasites. Flagellum and cell body length were measured with Image J as described previously [42]. Parasites were classified as metacyclic forms when the body length was < 14 μm, and flagella length ≥ 2 times body length. After measurements, the parasites were counted and expressed as percentage (%) of the metacyclic from each *Leishmania* species culture.

### Protein extraction and digestion

The samples were resuspended in 8 M urea, protease inhibitor cocktail (Sigma-Aldrich) and DNase I, followed by five freeze-thaw cycles for complete cell lysis. The extracted proteins were quantified by Qubit fluorimetric detection method. The solution was buffered with 50 mM NH_4_HCO_3_ (final concentration) and reduced by 10 mM DTT for 45 min incubation at 30°C. The proteins were subsequently alkylated by the addition of 40 mM iodoacetamide (IAA) and incubation for 30 min in the dark at room temperature, and the reaction quenched using 5 mM DTT. Porcine trypsin (1:50, w/w) (Promega) was added, and the mixture incubated overnight at 30°C. The reaction was stopped with 1% TFA and desalted using HLB hydrophilic–lipophilic-balanced solid phase extraction (Waters).

### Mass spectrometry analysis

Peptides were separated by nano-LC-MS/MS on Reprosil-Pur C18-AQ (3µm; Dr. Maisch GmbH, Germany) using Easy-LC nano-HPLC (Proxeon, Odense, Denmark) connected through a source of nanospray ions (Thermo Scientific) mass spectrometer Q Exactive™ HF hybrid quadrupole-Orbitrap (Thermo Scientific). The HPLC gradient was 0–34% solvent B (A, 0.1% formic acid; B, 90% ACN, 0.1% formic acid) at a flowrate of 300 nL/min. The data-dependent acquisition was done in positive mode. The 400-1600 m/z scan was recorded in Orbitrap at a resolving power of 120,000, target AGC of 3×10^6^ and maximum injection time of 100 ms. The MS/MS spectra of the 15 most abundant peptides were obtained by HCD (high energy C-trap dissociation) with acquisition in the orbitrap at a resolution of 30,000. The parameters for the acquisition of HCD were: isolation window: 1.2 Da; normalized collision energy: 28; Target AGC: 1×10^5^; dynamic exclusion: 30 s; intensity threshold: 3.3 x 10^4^ and maximum injection time of 150 ms. The Raw files have been submitted to the ProteomeXchange Consortium via the PRIDE [43] partner repository with the dataset identifier PXD044804.

### Protein identification, quantification and Bioinformatic analysis

LC-MS/MS raw files were imported into Proteome Discoverer version 2.5 for identification and label-free quantification (LFQ) of proteins and peptides. MS/MS spectra were searched against the combined Uniprot reference proteomes of *Leishmania mexicana* strain MHOM/GT/2001/U1103 (UP000007259)*, Leishmania braziliensis* (UP000007258) and *Leishmania infantum* database (UP000008153) (http://www.uniprot.org; downloaded June 5, 2022; 24173 entries), and a common contaminants protein database with a mass tolerance level of 4.5 ppm for MS and 0.05 Da for MS/MS. Enzyme specificity was set to trypsin with a maximum of two missed cleavages. Carbamidomethylation of cysteine (57.0215 Da) was set as a fixed modification. Oxidation of methionine (15.9949 Da), deamidation NQ (+ 0.9840 Da) and protein N-terminal acetylation (42.0105 Da) were selected as variable modifications. The ‘Feature Mapper’ node on Proteome Discoverer, which enables peptide identifications between samples based on their accurate mass, charge and retention time, was applied in the consensus workflow. All identifications were filtered in order to achieve a protein false discovery rate (FDR) of less than 1% [44-46]. Label free quantification (LFQ) of the proteins was calculated using the label free quantification algorithm embedded in the Minora feature detection node in the processing workflow. Contaminants were removed before further analyses using Perseus 2.0.7.0 computational platform [47]. For high confidence protein identification, top ranking proteins with the highest protein sequence coverage (master proteins) were chosen, and in addition, proteins identified with ≥ 2 peptides were selected for further analysis. Analysis of variance (ANOVA) was applied to identify the regulated proteins between the three *Leishmania* species and between each *Leishmania* species in the three different conditions using Benjamini-Hochberg-based FDR correction at a FDR<0.05. Differentially regulated proteins within the same *Leishmania* species between the conditions (growth stages and temperature) were determined using t-test statistical analysis with permutation-based FDR correction at a FDR <0.05, and presented as volcano plots. Clustering of the samples were evaluated according to the unsupervised hierarchical grouping of the regulated proteins after Z-score normalization. In addition, Gene Ontology (GO) analysis of statistically regulated proteins were mapped against protein expression values to identify the cellular components, biological processes and molecular functions differentially enriched in the three *Leishmania* species during parasite growth stages and temperature modulation. The GO analysis was performed in TriTrypDB for regulated proteins between species and between stages/conditions using *p*-value of 0.01 and 0.05 as a cutoff of enriched GO Slim terms, respectively.

### Viability and cell death assays

Parasite viability was checked before proteomic experiments. Parasites were washed twice with PBS, counted using Neubauer chamber, and plated (1×10^6^ per well) in M-199 medium and incubated for 5 min with 10 μg/mL propidium iodide (PI) (Sigma-Aldrich) to label the non-viable parasites. Parasites in the presence of 0.4% paraformaldehyde (PFA) were used as a positive control for PI labeling. The labeling was analyzed by fluorescence microscopy. The images were acquired in a DMI6000B / AF6000 (Leica) microscope coupled to a digital camera system (DFC365FX) and analyzed with Image J. In addition, the *Leishmania* species were cultured up to the 6^th^ day, washed twice with PBS, and counted. Approximately 2×10^6^ per well were plated in 96 well plates in M-199 medium supplemented, as described previously [41]. The plates were maintained at 25 and 34°C for 24 hours. The optical density was subsequently determined on a microtiter reader (BioTek) at reference wavelength of 690 nm and test wavelength of 595 nm. A standard curve containing different amounts of promastigotes (10^3^, 10^4^, 10^5^, 10^6^ and 10^7^ per well) was used for quantification.

### Thermotolerance Assays

To determine parasite resistance at different temperatures, 200 μL of cell suspensions containing 2 x 10^6^ of parasites were plated in 96-well plates and incubated at different temperatures: 8, 25, 34, 37 and 42°C. After 24 hours of incubation, viability assays were performed. For this, 30 μL of MTT [(3- [4,5-dimethyl-2-thiazolyl] -2,5-diphenyl-2H-tetrazolium bromide; 5 mg/mL; (Sigma-Aldrich)] was added in each well and the plate incubated for 3 hours at 25°C. Sodium dodecyl sulfate (50 μL of 10%) was used to solubilize the formazan crystals produced by the reaction, and the absorbance measured using a Microtiter plate reader (BioTek) at a reference wavelength of 690 nm and a test wavelength of 570 nm [41].

### Quantitative Real-Time PCR

To quantify the expression of HSP genes, the promastigote forms of the *Leishmania* spp., cultured for 6 days at 25°C were maintained at 25 and 34°C for 24 hours. The parasites were washed twice with PBS prior to RNA extraction and purification using the Quick-RNATM MiniPrep kit (Zymo Research, Irvine, CA, United States) according to manufacturer specifications. Quantification was carried out using a NanoDrop™ spectophotometer (Thermo Scientific), and the complementary deoxyribonucleic acid (cDNA) synthesis was performed using the SuperScript III Reverse Transcriptase kit (Invitrogen). Quantitative PCR reaction was performed in equipment StepOnePlusTM Real-Time PCR System (Applied Biosystems) using Maxima SYBR Green/ROX qPCR Master Mix (Thermo Scientific) according to the manufacturer’s instructions.

Oligonucleotide primers were used to amplify a portion of the different target molecules: HSP 83-1 cDNA forward, 5’-GATGACGACACTGAAGGACTAC-3’ and reverse, 5’-ACGACTCCAGCTTCTTCTTG-3’; HSP70 cDNA forward, 5’- ATCACGATCCAGAACGACAC-3’ and reverse, 5’-TGTCCTCCTCAGCAAACTTC-3’; Arginase cDNA (control) forward: 5’-GTGTCATACGACGTGGACACG-3’and reverse: 5’-CGACAAGACGACCGCACTCGGC-3’.

The reaction was incubated for 10 min at 95°C followed by 40 cycles of 15 s at 95°C, 30 s at 55°C and 30 s at 72°C. Fluorescence was detected at each annealing step. Technical triplicates were performed for each reaction, and no template controls (NTC) were included. The data were presented as relative quantification normalized by the level of the endogenous Arginase gene expression, calculated using 2^-ΔCt^ [48].

### Western blot analysis

Parasites were resuspended in Laemmli sample buffer, lysed by boiling at 95°C for 5 min, and the proteins separated on a 12% SDS-PAGE. The proteins were fixed and stained with Coomassie Brilliant Blue stain and destained for visualization. Lane intensities were analyzed using ImageJ software. Western blot analysis of HSP70, GP63 and α-Tubulin was performed by transferring proteins to nitrocellulose membranes (GE healthcare). The membranes were blocked, incubated overnight at 4°C with anti-HSP70 (1:10000, gently provided by Dr. Joachin Clos), anti-GP63 (1:5000; kindly provided by Prof. Rob McMaster) or anti-α-Tubulin (1:5000; Sigma 05829) primary antibodies, followed by incubation for 1 hour with anti-IgG mouse (1:10000; 741806) or anti-IgG chicken (1:160000; KPL 142406) HRP-conjugated secondary antibodies. Band intensities were analyzed using ImageJ software and the results were normalized based on protein intensities from total lysates previously analyzed by Coomassie Brilliant Blue stained lane intensities.

### Zymography assay

Parasites (1×10^8^) from *Leishmania* spp., were lysed by three freeze-thaw cycles followed by three cycles of sonication in lysis buffer (180 µL urea, 8M) complemented with 10 µL 10x protease inhibitor cocktail (Sigma-Aldrich) and 10 µL 10x phosphatase inhibitor cocktail (Sigma-Aldrich). Extracted proteins were quantified by the Qubit fluorometric method (Thermo Fisher), and 20 µg of proteins for each species analyzed by a SDS-PAGE gelatin zymography as previously described [49] with minor modifications. Briefly, a 10% SDS-PAGE gel was prepared by the addition of gelatin (Sigma) at 1 mg/mL. The samples were electrophoresed at a constant voltage of 125V, followed by the incubation of the gels in buffer 1 (50mM Tris-HCl pH 7.4, 2.5% Triton X-100) for 1 hour to remove SDS. Subsequently, the gels were washed three times with water and incubated at 37°C for 16 hours with buffer 2 (50 mM Tris-HCl, 10 mM CaCl_2_, 200 mM NaCl, 0.2% CHAPS), which increases gelatinase activity. The gels were stained with Coomassie Brilliant blue and gelatinase activity was measured. Images were obtained using the BioRad ChemiDoc XRS+ Molecular Imager with the Image Lab software and ImageJ using the reverse intensity area of the white bands. Normalization was based on the total protein concentration applied to the gel.

### Statistical analysis

Data were analyzed with Prism 6.0 (GraphPad Software, San Diego, CA). Statistical significance was determined by ANOVA One-way for data with Gaussian distribution and similar variation between groups. After test for normal distribution, non-Gaussian data were analyzed by Kruskal-Wallis or Dunn’s Multiple comparison test, as indicated. Results represent means ± standard deviation (SD) or standard error of the mean (SEM) as indicated. The number of independent experiments, technical and biological replicates are indicated in corresponding figure legends.

## RESULTS

The proteomes of *L. amazonensis (La), L. braziliensis (Lb)*, *and L. infantum (Lc)* were analyzed using a large-scale bottom-up mass spectrometry-based quantitative proteomics approach followed by statistical analysis and functional validation using orthogonal techniques (**Fig. 1**). After validation of the proliferation capacity and metacyclic differentiation (**Supplementary Fig. 1A, B**) for comparison, the *Leishmania* parasites were collected in three different conditions: exponential/logarithmic phase (Exp), stationary phase 1 (Sta25), and stationary phase 2 (Sta34) (**Fig. 1A)**. Parasites in the exponential/logarithmic phase were collected on day 3, while parasites at the stationary phase 1 were collected on day 7. Both conditions were maintained at 25°C. In order to study the effect of temperature changes and resembling parasite adaptation within the vertebrate host, we collected parasites in the stationary phase at condition 2, which was obtained from parasites on day 6, incubated at 34°C for 24 hours (**Fig. 1B**). Before proteomic analysis, the parasite viability was assessed by propidium iodide staining and fluorescence microscopy. Viability greater than 99% was detected (**Fig. 1C**). The label-free quantification algorithm (LFQ), embedded in the Minora feature detection node in PD, extracted the intensity of each identified peptide and inferred the relative normalized abundance of the proteins in the three species. These normalized intensities were used to compare the abundance of the proteins in the three species and the three conditions.

**Supplementary Fig. 1.**
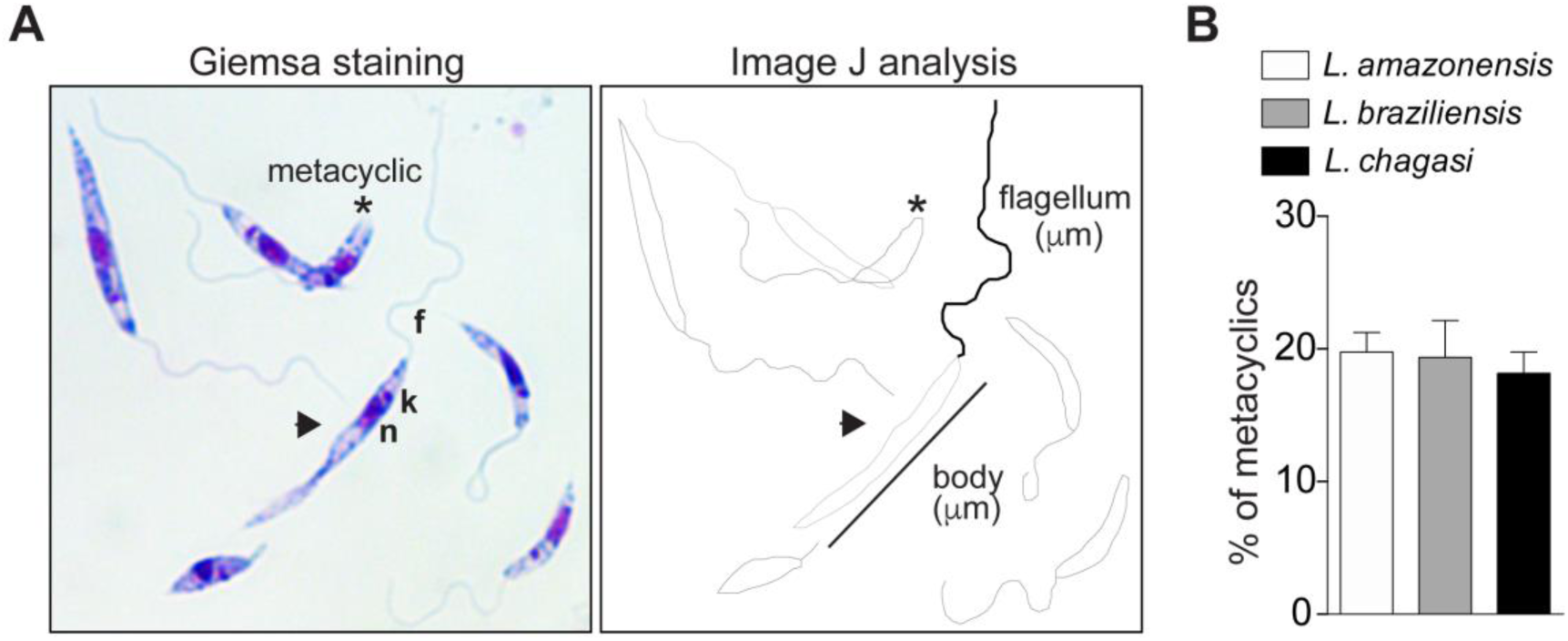
Confirmation of parasite culture synchronization from the different *Leishmania* species. Samples from each species were analyzed by comparing the number of metacyclic forms in the stationary phase of growth. (**A**) Representative images of the Giemsa staining from the samples were prepared to visualize different parasite structures such as flagellum (f), nucleus (n), and kinetoplast (k). To confirm the presence of metacyclic forms (*) from other parasite forms (arrowhead), further analysis of the different parasites was processed by Image J, analyzing the length of the flagellum and the body of the parasites. Metacyclic forms were classified when the body length was < 14 μm, and flagella length ≥ 2 times body length. (**B**) After measurements, the parasites were counted and expressed as % of the metacyclic from each *Leishmania* species culture. The representative graph corresponds to the mean ± SD of five biological replicate assays (one-way ANOVA; Kruskal-Wallis/Dunn’s multiple comparison tests).

A bottom-up large-scale mass spectrometry-based proteomics approach was applied to study the differential proteome of La, Lb and Lc. The three species were analysed in exponential/logarithmic (Exp), stationary 25°C (Sta25) and stationary 34°C (Sta34), with a total of 9350 proteins identified and quantified (**Supplementary Table 1**).

### Comparative proteome analysis of *L. amazonensis*, *L. braziliensis* and *L. infantum chagasi* in different growth phases and temperature stress

The first analysis focused on comparing the three species in each condition. Of the 9350 proteins identified and quantified in this analysis (**Supplementary Table 1a)**, 8890 proteins were identified with high confidence (**Supplementary Table 1b)**, while 460 proteins identified with medium confidence and were not considered for further analysis. For subsequent analysis, only proteins identified with 2 or more peptides were considered, totalling to 7047 proteins (**Supplementary Table 1c)**, which corresponded to a coverage of 88% of the *Leishmania* predicted proteome, considering ∼8000 protein-coding genes predicted and one proteoform per gene. Of these proteins, 2729 proteins are uncharacterized, corresponding to 39%. To further increase the identification confidence of the proteins analysed, only proteins identified in two replicates in at least one species/condition were considered. For the exponential phase, a total of 5885, 6061, and 5959 proteins were identified for La, Lb, and Lc species, respectively, and mapped with 7.9, 7.6 and 7.7 peptides per protein in La, Lb, and Lc, respectively (**Supplementary Table 2A, Fig. 2A**). For the stationary phase at 25°C, a total of 5598, 5929, and 5937 proteins were identified for the La, Lb, and Lc, respectively, and mapped with 7.9, 7.7, and 7.6 peptides per protein in La, Lb, and Lc, respectively (**Supplementary Table 2B, Fig. 2A**). For the stationary phase at 34°C, a total of 5754, 6014, and 5888 proteins were identified for the La, Lb, and Lc, respectively with 7.9, 7.6, and 7.7 peptides per protein in La, Lb, and Lc, respectively (**Supplementary Table 2C, Fig. 2A**). A total of 6878, 6655, and 6753 proteins were identified in the three species at the exponential, stationary at 25°C, and stationary at 34°C conditions, respectively (**Supplementary Table 1**, **Fig. 2A**). A higher number of unique proteins, 124 (representing 1.8% of the Exp proteins) was identified for the exponential phase, while 6 proteins (0.09%) and 20 proteins (0.29%) were identified exclusively in Sta25 and Sta34 phases, respectively (**Supplementary Table 2d, e, f**). More proteins were shared between the exponential and stationary phase at 34°C (165 proteins) compared to the two stationary phases (70 proteins) (**Fig. 2B**). The majority of the proteins identified as specific to the Exp stage are uncharacterized (40), while 6 ribosomal proteins, 3 kinases, 1 HSP60 (E9B155) and 1 leishmanolysin (A4H639) were also identified in this group. A higher number of proteins was detected in the exponential phase compared to the stationary phase at 25°C and stationary at 34°C in all species (**Fig. 2C, D, E**). Six protein were identified in the Sta25 phase, which included three uncharacterized proteins, a Putative glutamamyl carboxypeptidase, Ubiquitin-fold modifier 1 and GLF protein (**Supplementary Table 2e**). Sta34 specific proteins identified and quantified in this study were 20 (**Supplementary Table 2f**), including 8 uncharacterized proteins, dolichyl-diphosphooligosaccharide--protein glycotransferase, Putative tubulin tyrosine ligase, and a putative Proteasome regulatory non-ATP-ase subunit 8 among others. Negligible changes in the number of identified proteins was observed for the three *Leishmania* species from Sta25 to Sta34, with an increase in La (156 proteins) and Lb (85 proteins), corresponding to 2.8 and 1.4 percentage, respectively, while a 0.8% decrease was observed for Lc (49 proteins).

**Fig. 2.**
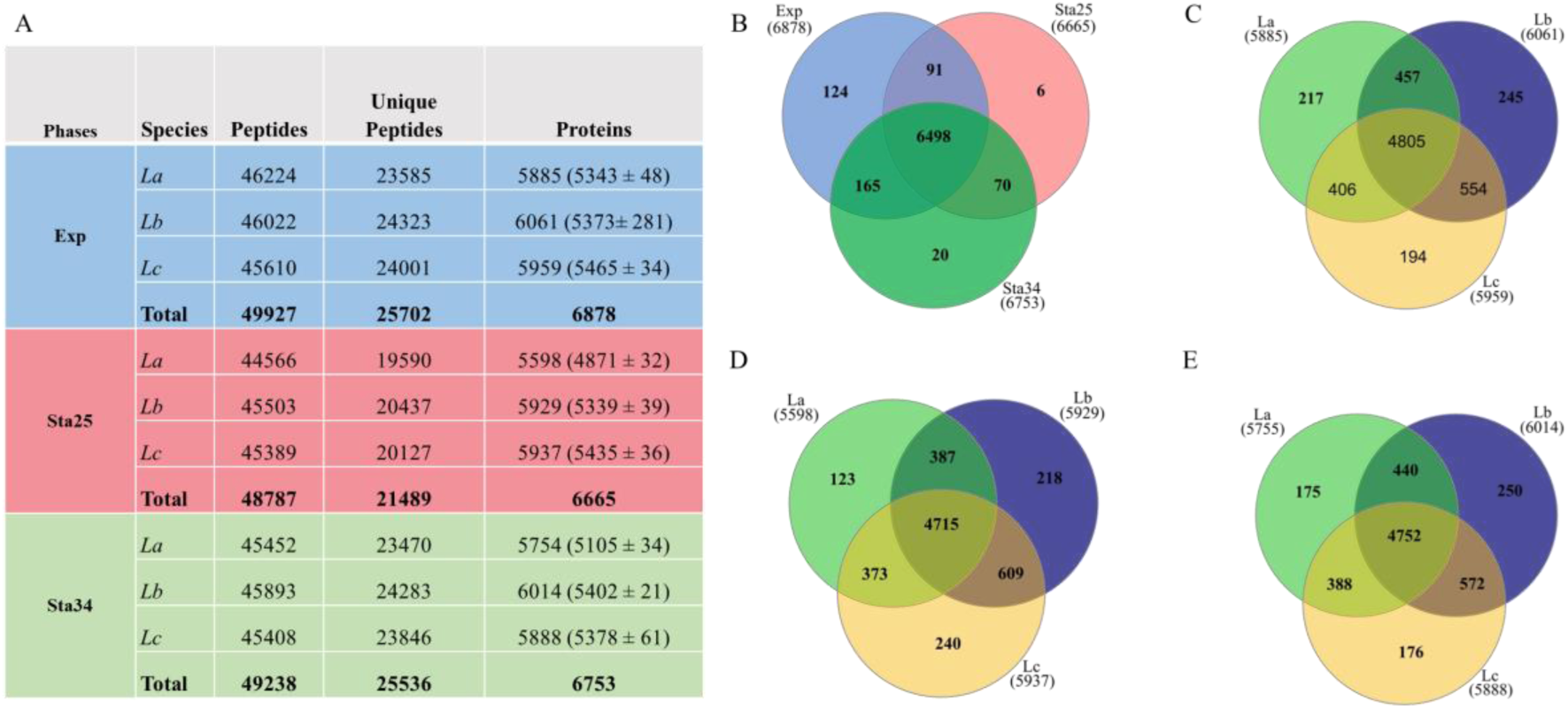
Proteins identified in *L. amazonensis* (La), *L. braziliensis* (Lb) and *L. infantum chagasi* (Lc) in the exponential and stationary phases at 25°C and 34°C. **A**) Number of peptides, unique peptides and proteins identified in the three species and conditions, followed by the mean and standard deviation of the three biological replicates. **B**) Venn diagram showing the number of shared and exclusive proteins between the three conditions: exponential (blue), stationary 25°C (pink) and stationary 34°C (green), and between the three species in **C**) exponential (Exp), **D**) stationary 25°C (Sta25) and **E**) stationary 34°C (Sta34), respectively.

Multivariate analysis of the total protein expression profile among the three species in the exponential phase revealed that La and Lc cluster closer together and well separated from Lb based on the first principal component which represents 46.2% of the total proteome variation in the dataset (**Supplementary Fig. 2A**). From a total of 7047 proteins identified in a minimum of two biological replicates in at least one species, 4129 proteins (58.59%) were regulated with Benjamini-Hochberg adjusted *p-*value < 0.05 among the three species at the exponential phase, as shown in the heatmap (**Supplementary Table 2G**), also highlighting the clustering of La and Lc separated from Lb based on Euclidean distance (**Fig. 3A**). Principal component analysis (PCA) of the total protein profile among the three *Leishmania* species in the stationary phase at 25°C highlighted the intermediate profile of *La* between the other two species (**Supplementary Fig. 2B**) based on the first component representing 49.8% of the total proteome variation. From a total of 6665 proteins identified in a minimum of two biological replicates in at least one species, 3831 (57.47%) proteins were differentially expressed between the three species at Sta25 condition with Benjamin-Hochberg adjusted *p*-value < 0.05 (**Fig. 3B** and **Supplementary Table 2H**). Lb presented 1272 proteins more expressed than the other two species (clusters f), of which 3.8% are represented by kinases, 2% ribosomal proteins, and 0.9% heat shock proteins. Clusters c and d were exclusively upregulated in La, with 627 and 425 proteins, respectively, while cluster a was specifically upregulated in Lc, with 649 proteins. Multivariate analysis of the total protein profile among the three *Leishmania* species in the stationary phase at 34°C showed a clearer separation between the subgenera *Leishmania* and *Viannia*, with La and Lc clade (*Leishmania Leishmania*) clustering closer together compared to the Lb clade (*Leishmania Viannia*) (**Supplementary Fig. 2C**). This profile was different from the stationary at 25°C, indicating a differential modulation of the proteome of the three *Leishmania* species upon a thermal shift. From a total of 6753 proteins identified and quantified in a minimum of two biological replicates in at least one species, 3994 (59.1%) proteins were differentially expressed between the three species with Benjamini-Hochberg adjusted *p*-value < 0.05 (**Fig. 3C and Supplementary Table 2I**). The most regulated proteins were involved in metabolic pathways. A total of 452 proteins were upregulated specifically in La (cluster a), 1041 proteins were upregulated exclusively in Lb (cluster f), while 659 proteins were upregulated only in Lc (cluster c). A total of 341 proteins were upregulated in Lb and Lc compared to La (cluster e), while 586 proteins were upregulated in La and Lb compared to Lc (cluster d). These data demonstrate the regulation of the *Leishmania* species-dependent proteome modulation with specific metabolic, cell motility, protein folding and signalling pathways activated.

**Fig. 3.**
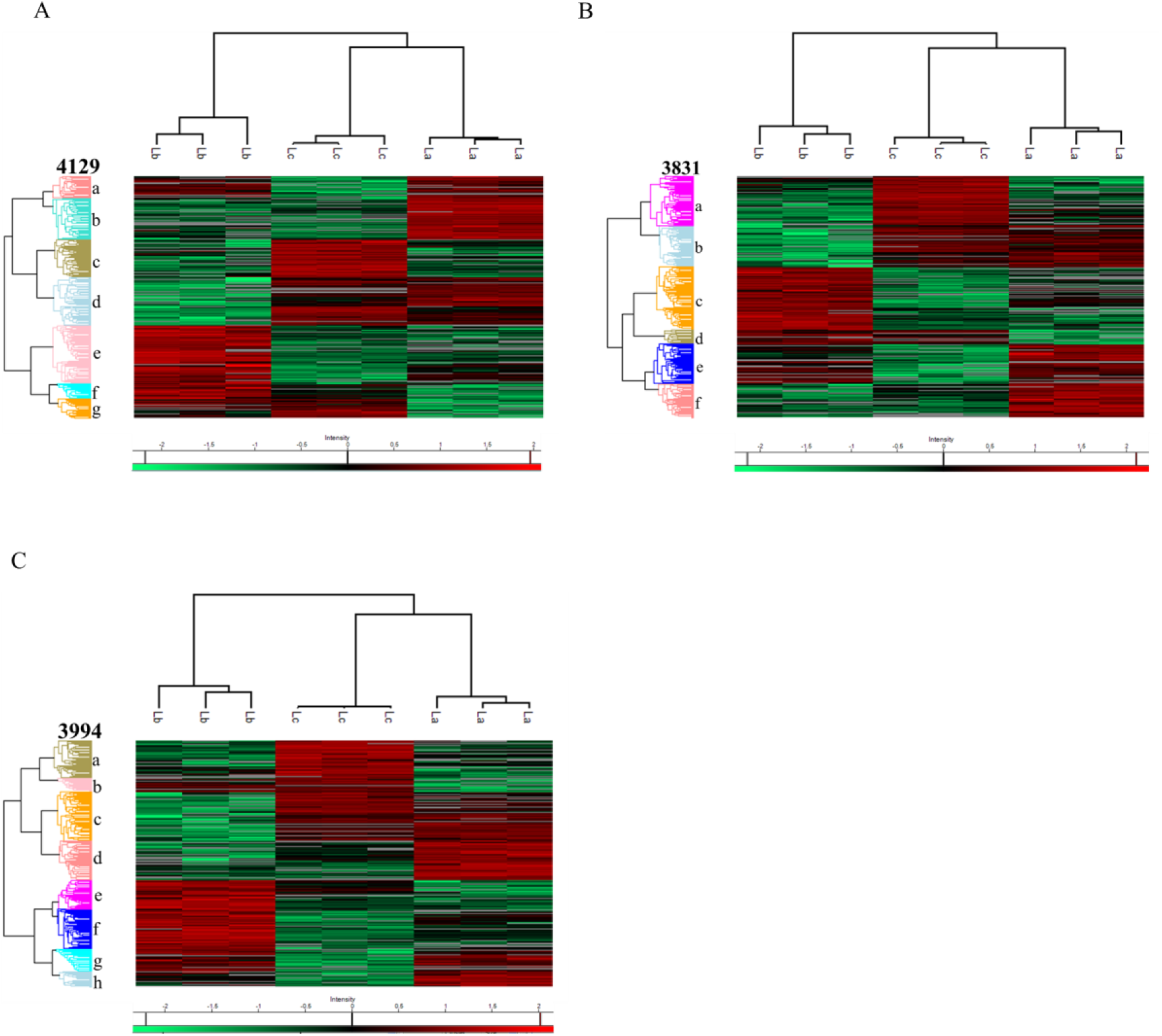
Comparison of identified and regulated proteins between *L. amazonensis* (La), *L. braziliensis* (Lb), and *L. infantum chagasi* (Lc). **A**) Exponential, **B**) Stationary 25°C and **C**) Stationary 34°C. Heatmap of the regulated proteins (ANOVA, p <0.05, with Benjamin-Hochberg correction), and Euclidean distance clustering considering at least 2 valid values per group/species. The most expressed proteins in a particular species are represented by the red colour, the least expressed by the green colour, absent in grey, and black, those that do not have a difference in the expression between two species.

**Supplementary Fig. 2.**
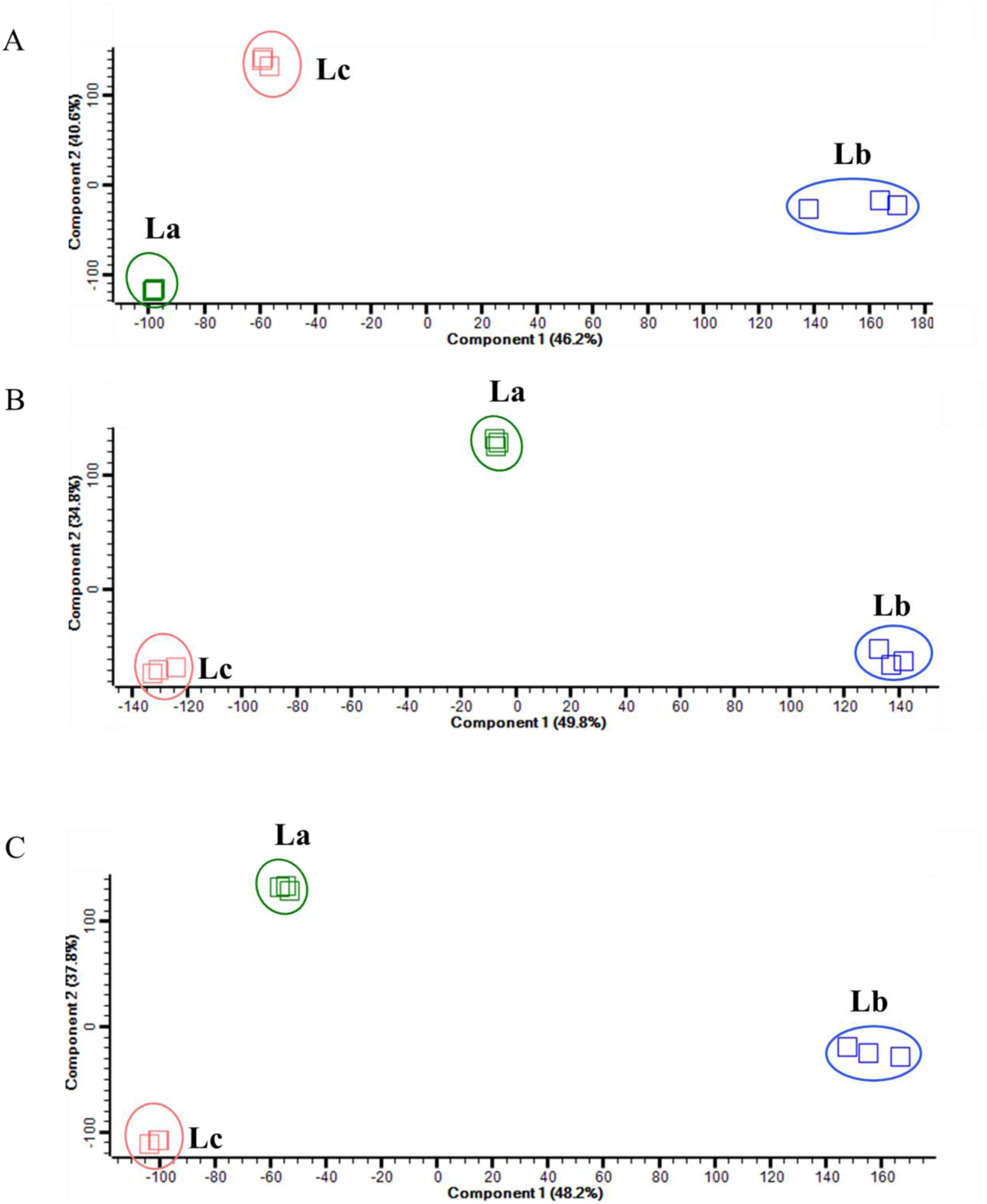
Multivariate analysis of the total proteomes of *L. amazonensis* (La), *L. braziliensis* (Lb) and *L. infantum* (Lc) in the three conditions. The principal component analysis (PCA) of La, Lb, and Lc in **A**) exponential, **B**) stationary at 25°C, and **C**) 34°C. Component 1 and 2 correspond to the total proteome variation and are expressed as a %.

To visualize proteins regulated between 2 species in the 3 conditions, a t-test with permutation-based FDR correction (FDR<0.05) was applied, and the regulated proteins were visualized by Volcano plots (**Supplementary Fig. 3).** Analysis of the regulated proteins at the exponential phase showed that 3133 proteins (45.6%) were regulated between La and Lb, 3382 proteins (49.2%) were regulated between La and Lc, and 3291 proteins (47.8%) were regulated between Lb and Lc (**Supplementary Fig. 3A, i-iii**). The accession numbers of the most abundant proteins are highlighted between each comparison in the volcano plots. Among the proteins expressed more abundantly at the Exp stage in La compared to Lb were heat shock proteins including Putative heat shock protein DNAJ (E9AZE9), Heat shock protein 83-1 (E9B3L2), Heat shock protein 70-related protein (E9AYA3); ribosomal proteins (A4IAU1) and proteins involved in cytoskeleton organization, such as Putative microtubule-associated protein (E9AKE4) and Tubulin alpha chain (E9AP62). At the Exp phase, most abundant proteins for Lb compared to La were proteins involved in metabolism, including Putative cystathione gamma lyase (A4HMZ0), Delta-12 fatty acid desaturase (A4HM35), Acetyl-coenzyme A synthetase (A4HCU1); and proteins involved in regulation of protein translation, including Putative translation elongation factor 1-beta (A4HAJ7) and Probable eukaryotic initiation factor 4A (Q25225). Comparison between La and Lc at Exp showed HSPs more abundant in La (Putative heat shock protein DNAJ (E9AZE9), Heat shock protein 83-1 (E9B3L2) and Putative heat-shock protein hsp70 (E9B099), Putative calpain-like cysteine peptidase (E9AUR8) and kinases including Regulatory subunit of protein kinase a-like protein (E9B573) and Putative serine/threonine-protein kinase (E9AXW8). Most abundant proteins for Lc in comparison to La at Exp phase included 60S ribosomal proteins (E9B6K7, A4IBL9, E9AQK4) and a Nucleoside diphosphate kinase (Q9GP00).

**Supplementary Fig. 3.**
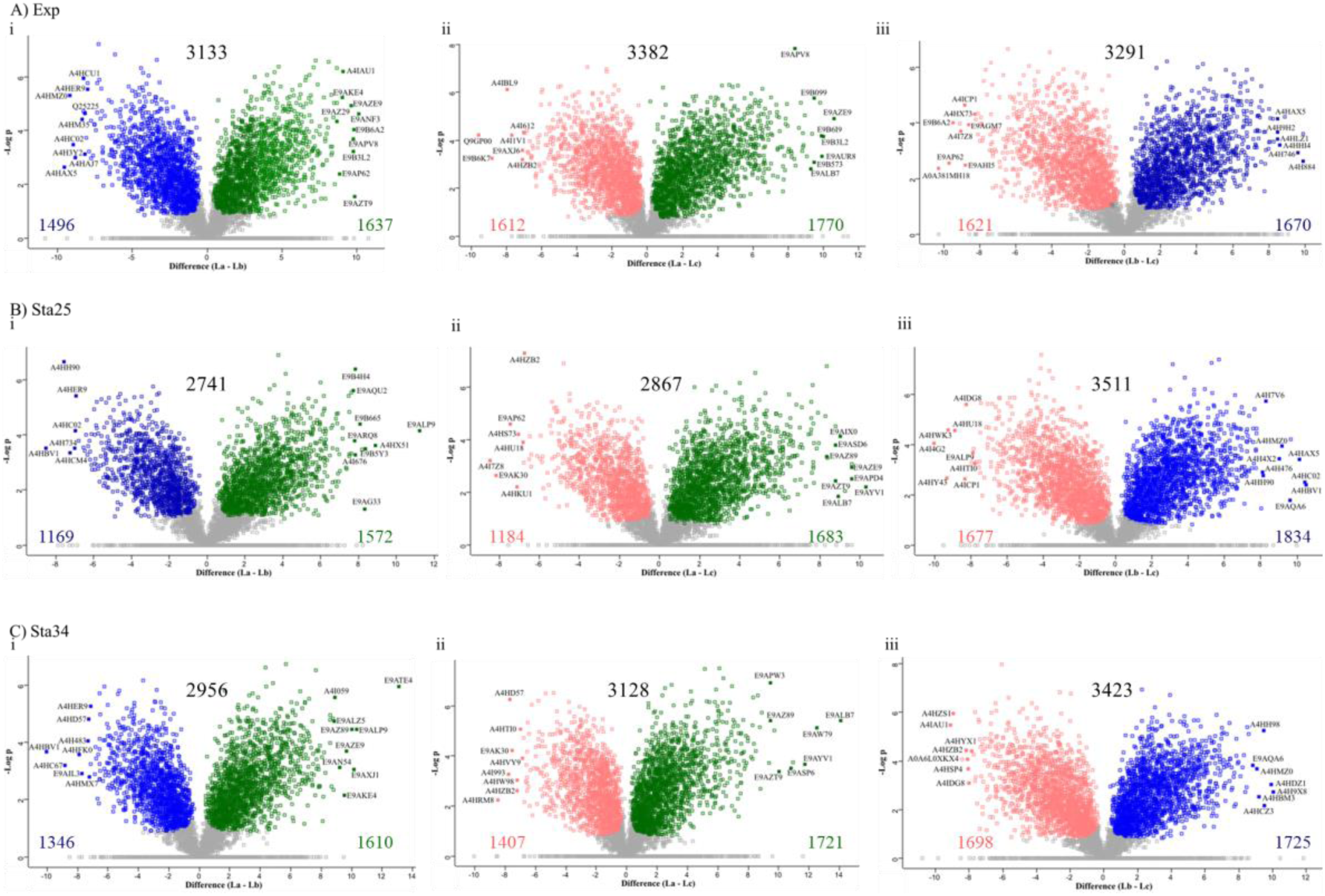
Volcano plots showing comparisons between two species for each condition. **The regulated proteins between A)** exponential (Exp), **B**) stationary at 25°C (Sta25), and **C**) stationary at 34°C (Sta34) between i) La and Lc, ii) La and Lc, and iii) Lb and Lc, respectively, are presented. Volcano plots performed with permutation-based FDR (FDR<0.05). La = green, Lb = blue, Lc = pink.

Regulation at the stationary phase at 25°C showed a different profile based on PCA analysis (**Supplementary Fig. 2B**), and the number of the regulated proteins between species decreased compared to Exp phase between La/Lb (2741) and La/Lc (2867), but increased between Lb/Lc (3511) (**Supplementary Fig. 3B, i-iii**). At 34°C, there was an increase in the number of regulated proteins between La/Lb (2965), and La/Lc (3128) compared to 25°C in the stationary phase, showing that at the higher temperature, mimicking the vertebrate skin, there is a greater protein modulation, differentiating the species from each other (**Supplementary Fig. 3C, i-iii**). However, a decrease from 3511 to 3423 regulated proteins was seen in Lb compared to Lc when shifting from 25 to 34°C during the stationary phase.

Differences in the proteome modulation while promastigotes were transitioning from the exponential to the stationary phases between the three *Leishmania* species were evidenced by gene ontology (GO) enrichment combined to protein abundances. At the exponential growth phase, proteins involved in biological processes such as metabolism of carbohydrate derivatives were enriched in *L. amazonensis*, and *L. chagasi* compared to *L. braziliensis*, while metabolism of sulphur compounds, microtubule based movement, protein folding and intracellular protein transport were enriched in La>Lb>Lc (**Supplementary Fig. 4A**). Proteins enriched in ribosome biogenesis were abundant in Lc compared to Lb. At Sta25, however, intracellular protein transport was enriched in Lb>Lc, and proteins involved in the metabolism of sulphur compounds were most abundant in Lb>La>Lc (**Supplementary Fig. 4D**). Unfolded protein folding, oxidoreductase activity, cytoskeletal protein binding and were among the molecular functions enriched in both Exp and Sta25 (**Supplementary Fig. 4B, E**). Proteins involved in oxidoreductase activity were abundant in La>Lb>Lc in both conditions, while proteins enriched in translation Lb>Lc>La. Cytoskeletal protein binding was enriched in La>Lb>Lc at the Exp phase, while at the Sta25 phase the proteins involved in the biological process were most expressed in Lb>Lc, but not enriched in La. In both growth conditions, enriched cellular components showed the upregulation of proteins in the cilium in *L. L. amazonensis* and *L. chagasi* compared to *L. braziliensis*, while proteins pertaining to the cytoskeleton were upregulated in La>Lb>Lc at the Exp phase, and at the Sta25 phase the cytoskeleton proteins were least abundant in Lb (**Supplementary Fig. 4C, F**). In addition, peroxisome proteins were upregulated in *L. braziliensis* compared to *L. amazonensis* and *L. chagasi* in both growth conditions.

**Supplementary Fig. 4.**
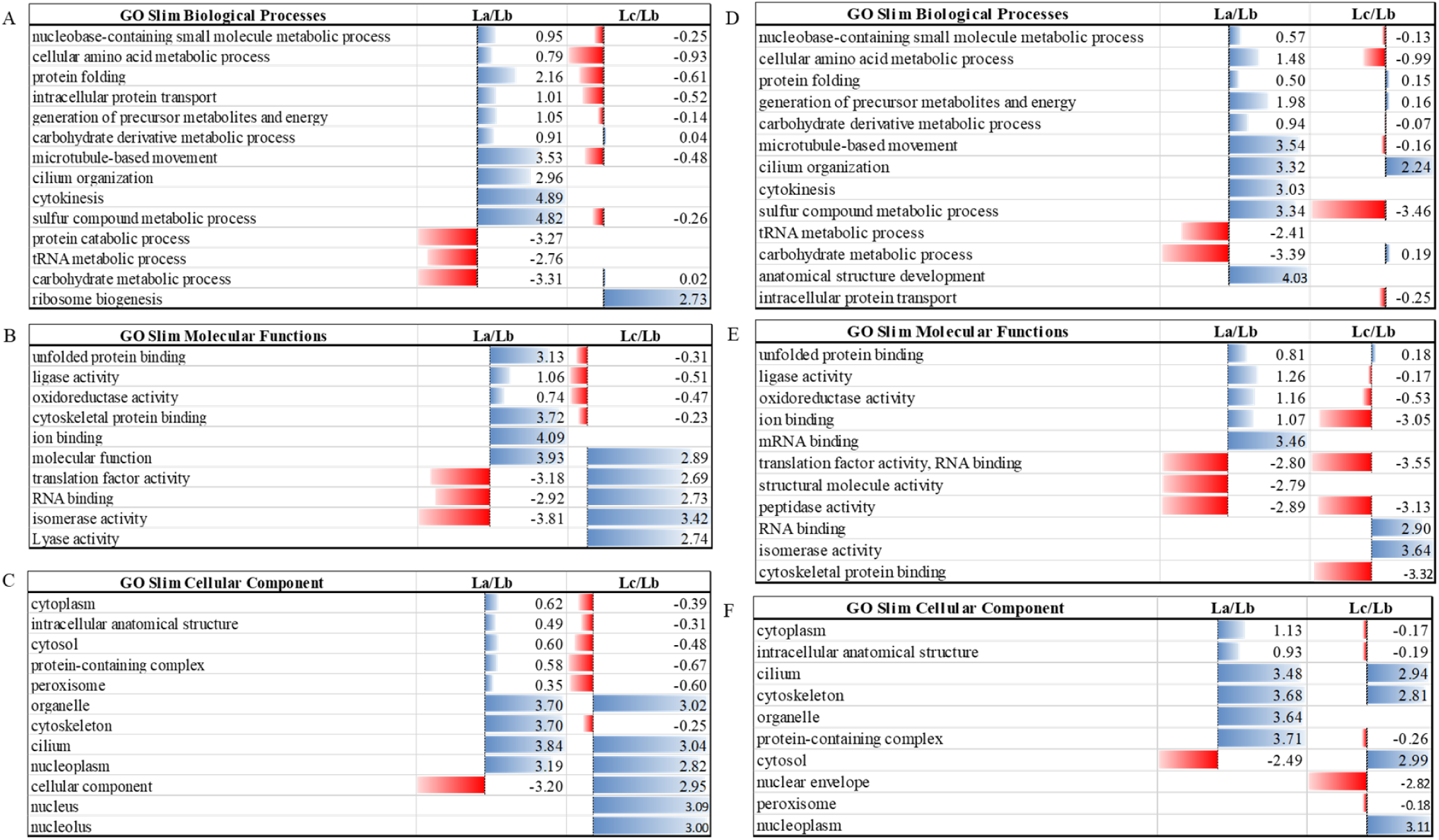
Gene ontology analysis combined with quantitative proteomics data of differentially expressed proteins between *L. amazonensis*, *L. braziliensis* and *L. infantum* at the exponential and stationary growth phases. The GO slim analysis shows the enriched biological processes, molecular functions, and cellular compartments at the exponential (A, B, C), and stationary at 25°C (D, E, F) growth phases, respectively. Blue bars represent proteins upregulated in *L. amazonensis* and *L. infantum chagasi*, and red bars represent proteins upregulated in *L. braziliensis*.

The proteome-wide rewiring during temperature shift from 25 to 34°C was also evidenced by GO analysis, highlighting species-specific enrichment of proteins involved in different biological processes, molecular functions, and cellular compartments between the three *Leishmania* species (**Supplementary Fig. 5**). Biological processes involving tRNA and carbohydrate metabolism, in addition to intracellular protein transport were abundant in *L. braziliensis* compared to *L. amazonensis* and *L. infantum*, while proteins involved in microtubule based moved were least enriched in Lb but abundant in La and Lc (**Supplementary Fig. 5A**). However, proteins involved in tRNA metabolic process, carbohydrate metabolic process and intra-cellular protein transport were most abundant in Lb compared to La and Lc at Sta34, while sulphur compound metabolic process was enriched in La and Lb and least in Lc. Over-representation of proteins with molecular function GO slim terms relating to unfolded protein binding, ligase activity, oxidoreductase activity and ion binding activity were least abundant in Lc and abundant in La>Lb, while translation activity and isomerase activity were enriched in Lb compared to La and Lc (**Supplementary Fig. 5B**). Interestingly, peptidase activity was enriched only in Sta25 and Sta34, where Lb and La had most abundant proteins involved in this GO term at the Sta25 and Sta34 conditions, respectively. Cellular components related to cilium, cytoskeleton and cytosol were enriched in La and Lc compared to Lb, while the cytoplasm, peroxisome and protein-containing complex were least enriched in Lc compared to La and Lb (**Supplementary Fig. 5C**).

**Supplementary Fig. 5.**
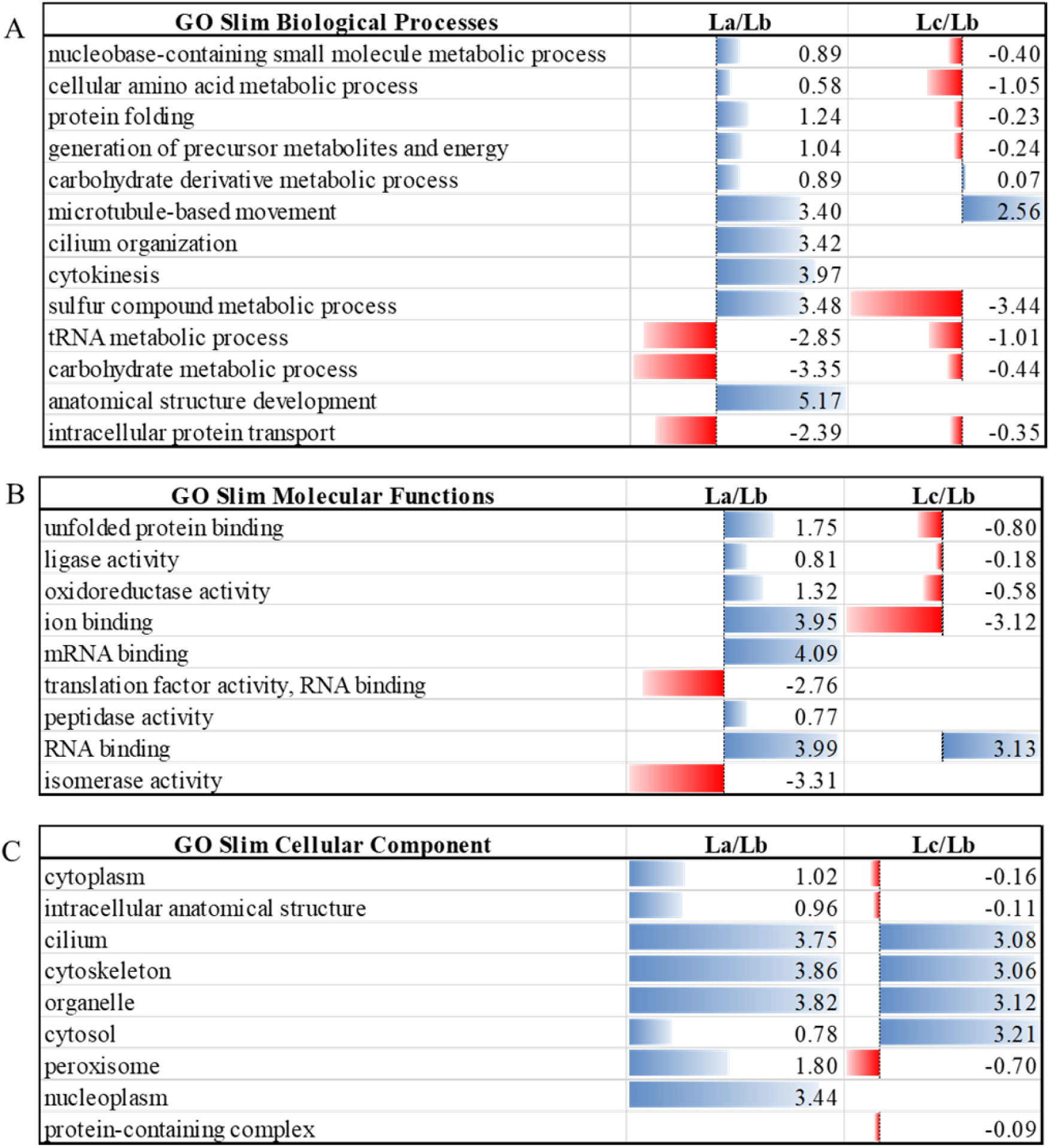
Gene ontology analysis combined with quantitative proteomics data of differentially expressed proteins between *L. amazonensis* (La), *L. braziliensis* (Lb), and *L. infantum* (Lc) during temperature transition from 25 and 34°C. GO slim terms representing (**A**) Biological processes, (**B**) Molecular functions, and (**C**) Cellular compartments. Blue bars represent proteins upregulated in *L. amazonensis* and *L. chagasi*, and red bars represent proteins upregulated in *L. braziliensis*.

Since some of the main biological processes and molecular functions regulated between the three species and upon growth phase transition were involved in cytoskeleton remodeling, proteolytic activity, and protein binding, we validated by western blotting the expression of some members of these classes, such as α-tubulin, gp63, and HSP70. Each band was quantified and normalized for the total protein loading measured by Coomassie staining. The expression of α-tubulin was higher in Lb compared to La (**Fig. 4A**). An antibody raised against pan α-tubulin did not give any signal for Lc, and it was not possible to report any quantification (**Fig. 4A**). The mass spectrometry-based α-tubulin quantification between the three species allowed the detection of two α-tubulin proteins, E9AP62 (LMXM_13_0280) and A4H727 (LBRM_13_0190), which were regulated in all three conditions. E9AP62 was downregulated in Lb at Exp and Sta34 conditions, while at Sta25 it was downregulated in La compared to La. α-tubulin A4H727 was most expressed in Lb at Sta25 and Sta34 and Lc compared to La (Lb>Lc>La), while at the Exp phase it was least abundant in Lc (La=Lb>Lc) (**Supplementary Fig. 6A**).

**Fig. 4.**
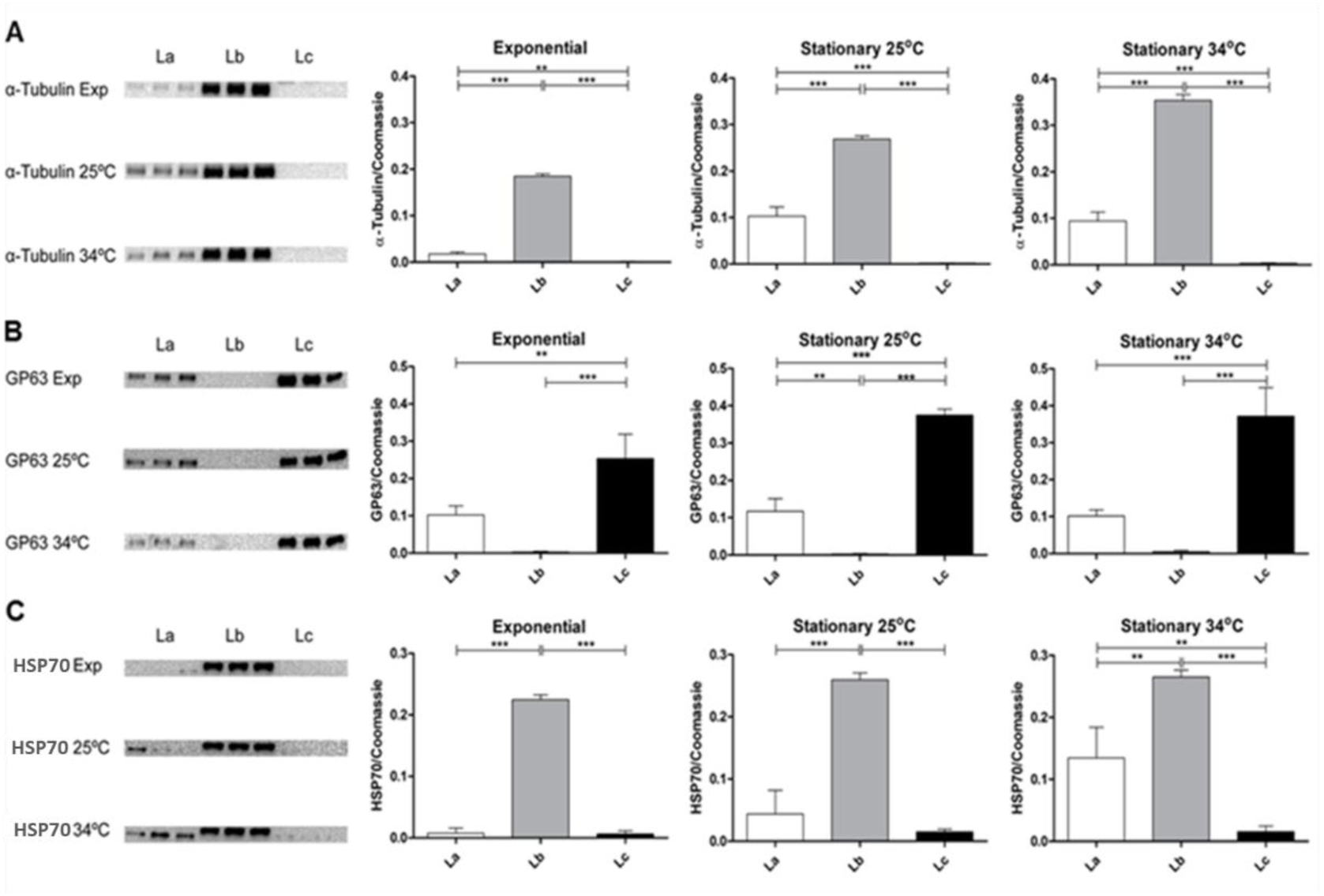
Comparative analysis of α-tubulin (A), GP63 (B) and HSP70 (C) abundances in the total protein extracts of promastigotes from exponential (Exp) phase, and stationary phases (Sta25 and Sta34) temperature shift. Western blot was performed with biological triplicates for *L. amazonensis* (La), *L. braziliensis* (Lb) and *L. infantum* (Lc). Data represented as mean ± standard deviation. Values were considered significantly different for p-values<0.05 when analysed by one-way ANOVA.

**Supplementary Fig. 6.**
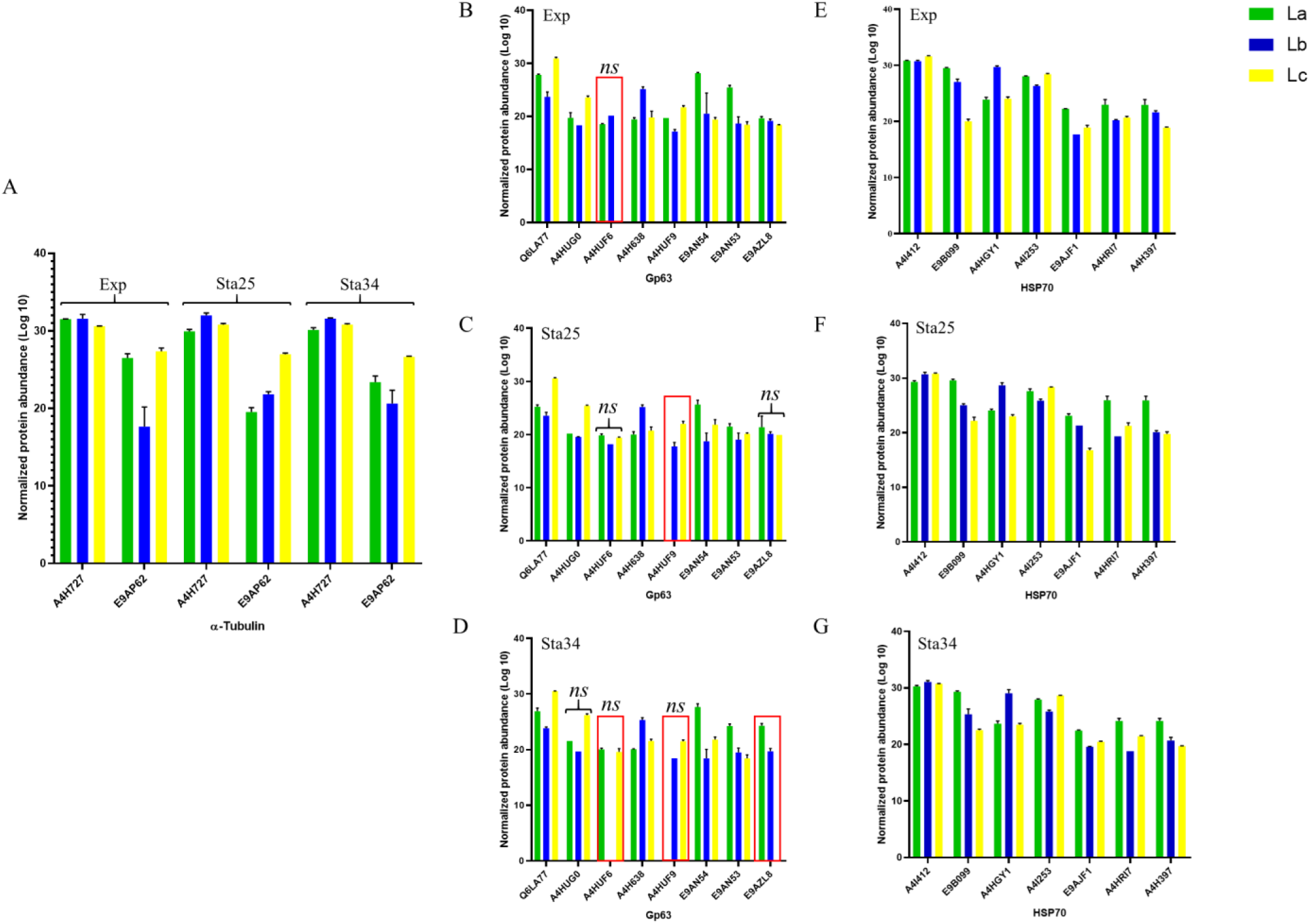
Identified and quantified α-tubulin, gp63, and HSP proteins in the three conditions chosen for validation by orthogonal techniques. The identification of **A**) α-tubulin from the different *Leishmania* species in the three conditions is shown. Identified and quantified gp63 in **B**) exponential (Exp), **C**) stationary 25°C (Sta25), and **D**) stationary 34°C (Sta34) conditions are shown. Identified and quantified HSP70 proteins in **E**) Exp, **F**) Sta25, and **G**) Sta34. All the proteins were regulated based on ANOVA with Benjamin-Hochberg correction (p <0.05), except the proteins marked by *ns,* while the proteins highlighted with the red box were proteins which were not identified in all three species in the different conditions.

The expression of gp63 was higher in Lc than La in the three experimental conditions (**Fig. 4B**). Interestingly, no signal of gp63 was observed in Lb. Mass spectrometry-based proteomics analysis allowed the detection of eight gp63 proteins: A4H638 (LBRM_10_0590), Q6LA77 (GP63-2), A4HUF6 (GP63-4), A4HUF9 (GP63-3), A4HUG0 (GP63-3), E9AZL8 (LMXM_28_0570), E9AN53 (LMXM_10_0460) and E9AN54 (LMXM_10_0390). In the exponential phase, all gp63 proteins were regulated except A4HUF6, which was identified in La and Lb but not in Lc (**Supplementary Fig. 6B**). Three gp63 proteins were upregulated in La (E9AN53, E9AN54 and E9AZL8), three were upregulated in Lc (Q6LA77, A4HUF9, and A4HUG0), while one (A4H638) was upregulated in Lb. (**Supplementary Fig. 6B, D)**. In the stationary phase at 25°C, two gp63 proteins were not regulated (A4HUF6 and E9AZL8), while A4HUF9 was found to be regulated in Lc and Lb (Lc>Lb) but not identified in La (**Supplementary Fig. 6C**). In the stationary phase at 34°C, it is possible to note more gp63 without regulation in their abundances (A4HUG0, A4HUF6 and A4HUF9) as well as more gp63 missing in certain species such as A4HUF9 for La, A4HUF6 for Lb and and E9AZL8 for Lc (**Supplementary Fig. 6D**).

The expression of HSP70 was higher in Lb than La and Lc in the three experimental conditions (**Fig. 4C**). Mass spectrometry-based quantitative proteomics revealed the regulation of seven HSP70 proteins, which were identified in all three *Leishmania* species and regulated in all three conditions. In the exponential phase, La had more HSP proteins with higher abundances (E9B099; E9AJF1, A4HRI7 and A4H397), only one was most abundant for Lb; A4HGY1 (LBRM_28_2990), while two were upregulated in Lc; A4I412 (LINJ_28_2960) and A4I253 (HSP70.4) (**Supplementary Fig. 6E**). A4HGY1 was upregulated in Lb compared to La and Lc (Lb>La=Lc). A4I253 was upregulated in Lc and La than Lb (Lc = La> Lb). E9B099 was upregulated in La compared to Lb and Lc (La>Lb>Lc). In the stationary phase at 25°C (**Supplementary Fig. 6F**), A4HGY1 presented the same profile as in the exponential phase, with higher expression in Lb compared to La and Lc (Lb>La=Lc). Moreover, the same HSP with highest abundances in La at the exponential phase were also identified in Sta25 with higher expression compared to Lb and Lc (E9B099; E9AJF1, A4HRI7 and A4H397), while A4I412 and A4I253 presented higher expression in Lc compared to La and Lb. In the stationary phase at 34°C (**Supplementary Fig. 6G**) A4HGY1 presented higher expression in Lb compared to the other two species (Lb>Lc=La), and E9B099 and A4H397 were least expressed in Lc compared to La and Lb (La>Lb>Lc). The regulation of A4HGY1 HSP70 protein confirmed the western blot data, also showing the differential regulation of HSP70 proteoforms.

The proteolytic activity was evaluated by zymography using collagen as a substrate, showing a different pattern between the three species and conditions (**Supplementary Fig. 7A**). Quantitation of the total proteolytic activity in the exponential phase revealed a higher activity for La followed by Lb and Lc (**Supplementary Fig. 7B**). In the stationary phase at 25°C, higher activity was detected in La followed by Lc and least in Lb. The same pattern as Sta25 was observed for the stationary phase at 34°C. The quantification of the proteolytic activity of the 63kDa band showed higher activity in La > Lc > Lb at the exponential phase. At the same time, at both Sta25 and Sta34 conditions, Lc had the highest activity, followed by La and minor activity recorded in Lb (**Supplementary Fig. 7C**).

**Supplementary Fig. 7.**
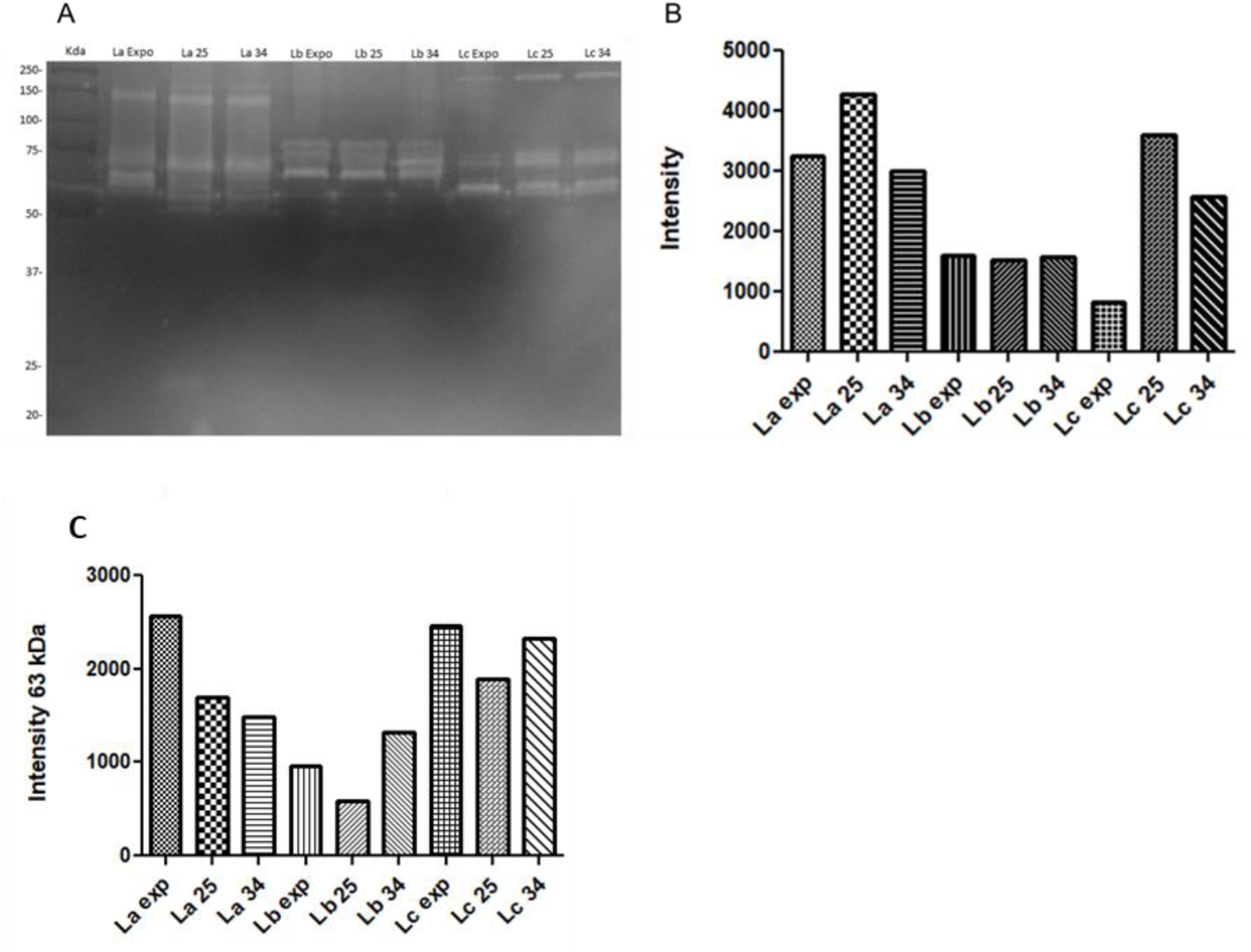
Proteolytic activity of *L. amazonensis*, *L. braziliensis*, and *L. infantum* in exponential (Exp), stationary 25°C (Sta25) and stationary 34°C (Sta34) conditions. **A**) Degradation of collagen by the protein extracts run on SDS-PAGE. **B**) Total quantification of proteolytic activity expressed as intensity of degradation bands. **C**) Total quantification of the proteolytic activity of the 63kDa band.

### Comparative proteome analysis of different growth stages and temperature in each *Leishmania* species

The second proteomic analysis was focused on comparing the three conditions for each *Leishmania* species. The expression profiles of the regulated proteins were visualized using heatmap clustering, showing the separation of the exponential phase from the two stationary phases at 25°C and 34°C in all three *Leishmania* species (**Figure 5**). A total of 2432 proteins were regulated in La, comparing the three conditions (**Fig. 5A, Supplementary Table 4A**). Multivariate analysis using the identified La proteins separated the exponential phase from the other stationary conditions based on the first principal component, representing 45.2% of the total proteome variability (**Supplementary Fig. 8A**). A total of 1380 proteins were regulated between the two stationary phases (**Supplementary Fig. 8B)**, 595 upregulated in Sta34, of which 8 and 7 were heat shock proteins and paraflagellar proteins, respectively. A total of 785 were upregulated in Sta25 with 7 HSPs and 7 paraflagellar proteins A total of 579 proteins were regulated in Lb in the three conditions (**Fig. 5B, Supplementary Table 4B**). PCA analysis of Lb revealed differential proteome modulation, showing a clear separation in the three conditions, with the exponential phase separated from the two stationary phases based on the first principal component representing 36.1% of the total proteome variation (**Supplementary Fig. 8C**). However, only 9 proteins were regulated between the two stationary phases (Sta25 and Sta34), were three (E9ARW6, A4HAF3, A4HHK5) and six (A4HHS5, A4HHS0, E9B4H4, A4HZI3, E9B5Y3 and A4IDB4) proteins were upregulated in Sta25 and Sta34, respectively (**Supplementary Fig. 8D).** A total of 2582 regulated proteins were identified in *L. chagasi* (**Fig. 5C, Supplementary Table 4C**). From the 651 regulated proteins between two stationary phases, 278 are present in Sta34, with 5 heat shock proteins, while 373 were present in Sta25, of which 4 were HSPs and 3 were paraflagellar proteins.

**Fig. 5.**
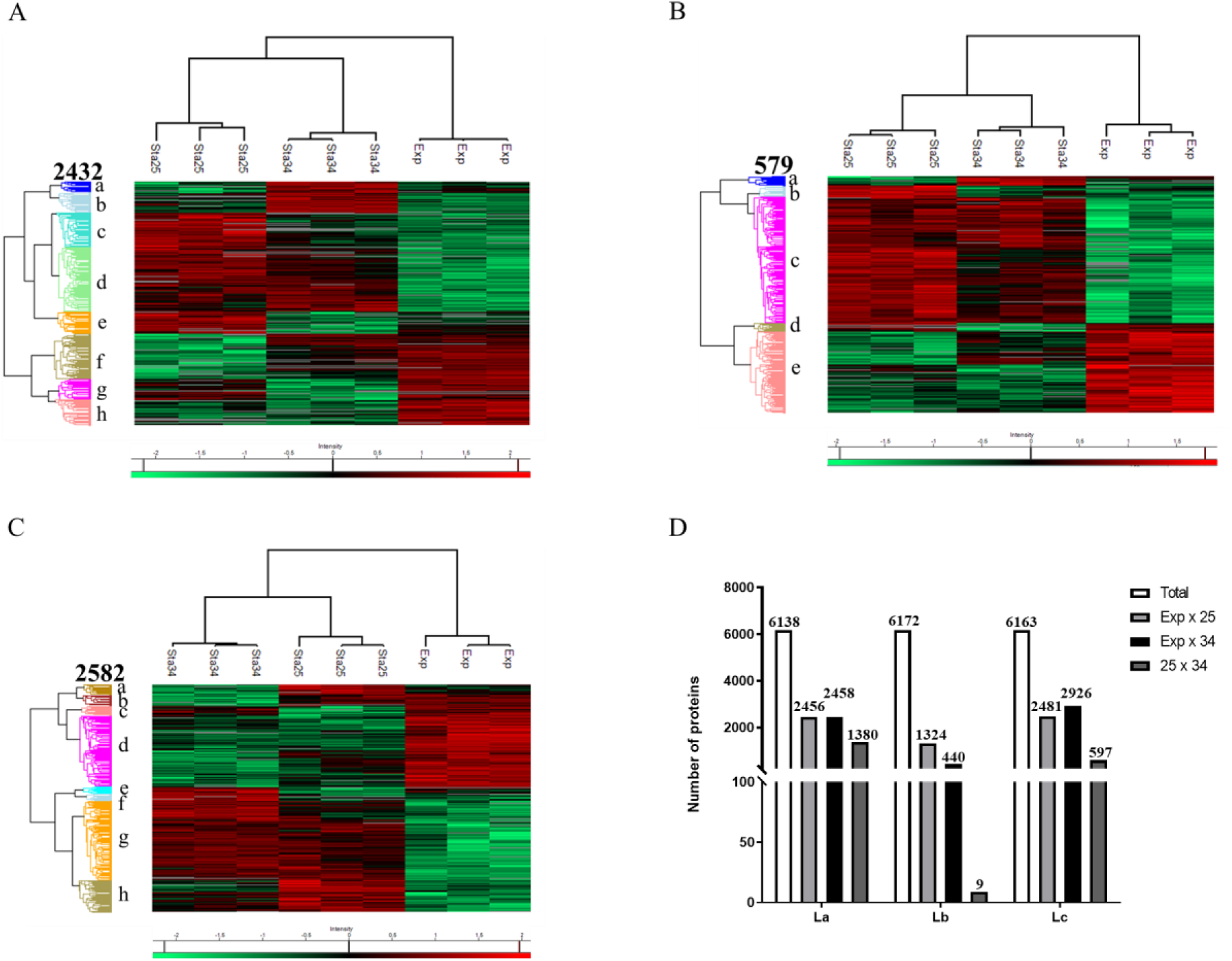
Comparison of identified and regulated proteins within growth phases and temperature shift. Heatmaps showing the regulated proteins (ANOVA, p <0.05, with Benjamin-Hochberg correction) using at least 2 valid values per group/species for **A**) *L. amazonensis*, **B**) *L. braziliensis,* and **C**) *L. infantum.* Red color represents the most expressed proteins in each species, the least expressed by the green color, absent in gray and black for proteins that do not present a difference in expression. **D**) Comparison of identified and regulated proteins in the three *Leishmania* species within exponential and the two stationary phases. The regulated proteins were based on Volcano plots based on permutation-based FDR (FDR <0.05).

**Supplementary Fig. 8.**
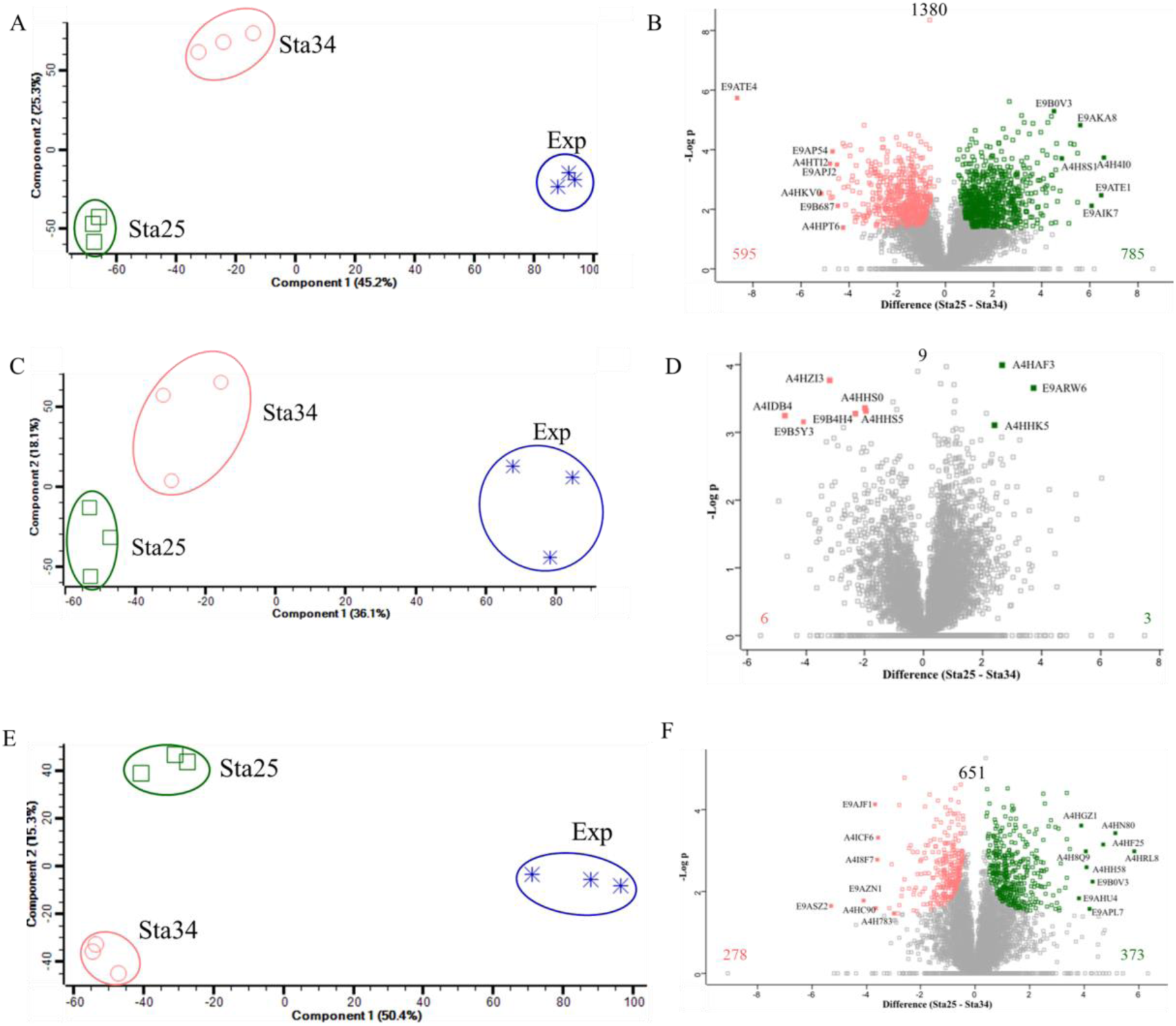
The *Leishmania* species’ proteomic profiles in exponential and stationary phases at 25°C and 34°C. The PCA clustering profiles in the first and second components and Volcano plots of La (**A, B**), Lb (C, D), and Lc (**E, F**) show total proteome variability at the Expo, Sta25 and Sta34 conditions, and significantly expressed proteins between Sta25 (green) and Sta34 (pink) conditions (FDR <0.05).

A total of 6138 proteins were identified in La, presenting differences between the three conditions (**Fig. 5C**). A total of 6163 proteins were identified in Lc, being 2926 (47.5%) regulated between exponential and Sta34 (**Fig. 5C**), showing a higher difference between these two conditions. A total of 6172 proteins were identified in Lb, of which 1324 proteins (21.5%) were regulated between the exponential and Sta25 (**Fig. 5C**). These results confirm a species-specific regulation in the transition from the exponential to the stationary phase.

Gene ontology analysis showed the proteome modulation while transitioning from the exponential to the stationary phases of the promastigotes in the three *Leishmania* species. The upregulation of proteins involved in biological processes, including protein folding and cellular amino acid metabolic processes were shown in the three *Leishmania* species at the exponential growth phase, while microtubule-based movement, cilium organization and intracellular protein transport were enriched at the stationary phase at 25°C in one or all Leishmania species **(Supplementary Fig. 9A)**. Cell mortality was enriched in Lb at the stationary growth phase, while proteins involved in carbohydrate metabolic process were abundant at the stationary phase in La but contrary in Lc they were they were abundant in the exponential phase. Contrary, protein catabolic process was upregulated in La at the exponential phase but in Lc this process was enriched at the stationary growth phase. Molecular functions involved in oxidoreductase activity and RNA binding were upregulated in the exponential growth phase in La and Lc, while peptidase activity, ion binding, and unfolded protein binding were only enriched in La at the exponential growth phase **(Supplementary Fig. 9B)**. Molecular functions related to structural molecule activity and structural constituent of ribosomes were upregulated exclusively in Lb during the exponential growth phase. Cellular components including intracellular anatomical structure and peroxisomes were upregulated in all species during the exponential growth phase, while the cytoskeleton was enriched at the stationary phase in all three species, with the highest enrichment in Lb > La > Lc (**Supplementary Fig. 9C**). The cytoplasm was enriched at the exponential phase in La and Lc, but at the stationary phase in Lb. Cilium was upregulated in the stationary phase in La and in Lc.

**Supplementary Fig. 9.**
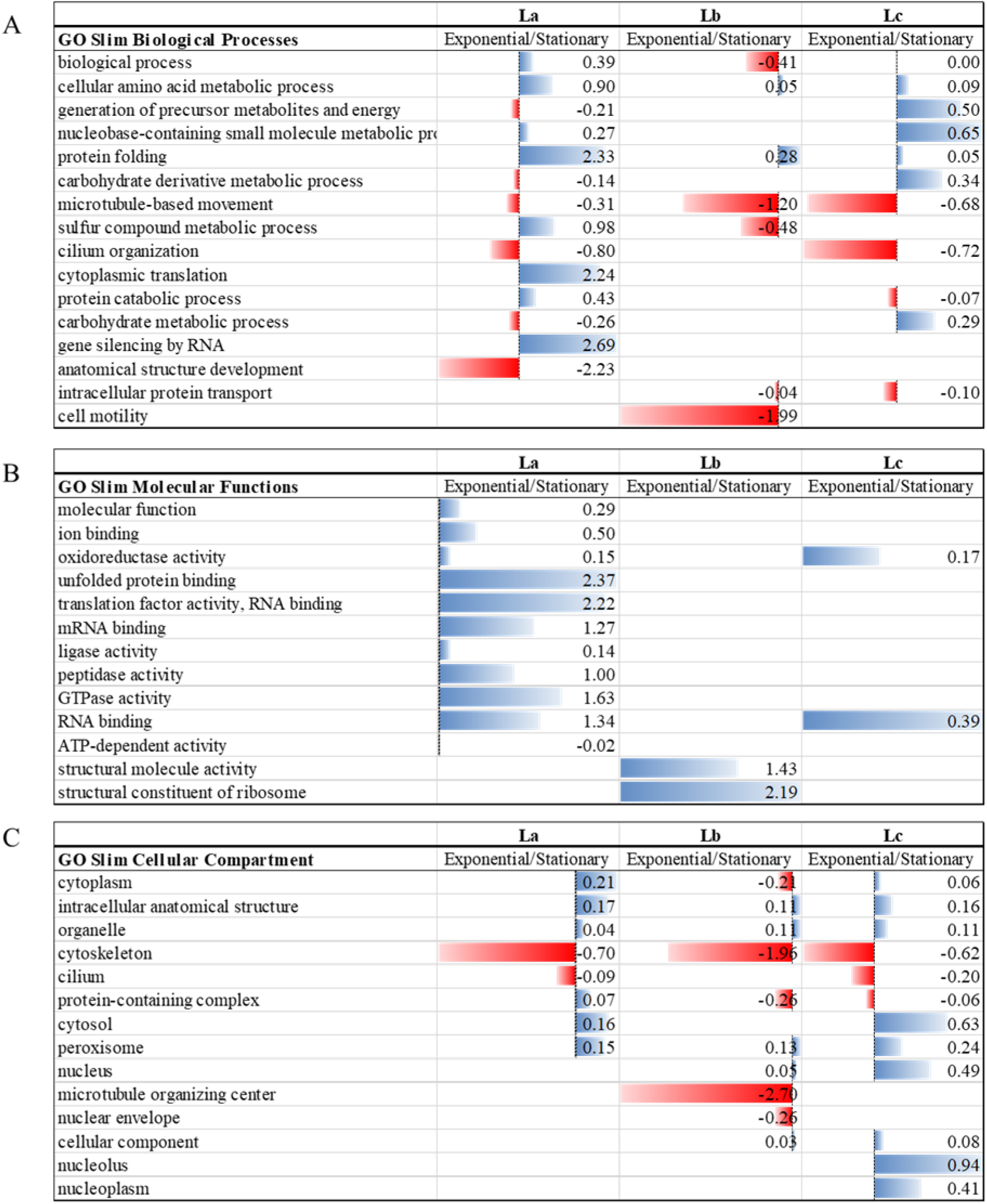
Gene ontology analysis combined with quantitative proteomics data of differentially expressed proteins during the transition from exponential to stationary growth phase for *L. amazonensis*, *L. braziliensis*, and *L. infantum*. (**A**) GO slim terms showing the Biological processes, (B) Molecular functions, and (**C**) Cellular compartments enriched in each *Leishmania* species. Blue bars represent proteins upregulated in the exponential phase, while red bars represent proteins upregulated in the stationary phase at 25°C.

The proteome-wide rewiring during heat stress from 25°C to 34°C was species-specific for La, Lb and Lc, as illustrated by GO slim analysis **(Supplementary Fig. 10)**. Notably, only nine (9) proteins were regulated during temperature shift from 25°C to 34°C in Lb, showing an enrichment of GO term related to signaling with higher abundance in Sta34 compared to Sta25. Enrichment of proteins from biological processes involving protein folding and nucleobase-containing small molecule metabolic processes were enriched at 34°C compared to 25°C in La but in Lc these processes were enriched at 25°C (**Supplementary Fig. 10A**). Microtubule-based movement and cilium organization were enriched in La at 25°C, reinforcing the species-specific modulation during temperature shift from 25° to 34°C. In addition, the cilium organization and microtubule-based movement were enriched at 25°C in *Leishmania* amazonensis, while protein folding was enriched in La at Sta34 but at Sta25 in Lc. The upregulation of proteins involved in molecular functions including RNA binding and translation factor activity were enriched at Sta25, while unfolded protein binding, oxidoreductase activity and peptidase activity were enriched during heat stress in La. For Lc, proteins involved in cytoskeletal protein binding and glycosyltransferase activity were enriched in Sta25 compared to Sta34. The cellular components upregulated in both strains at 34°C included the cytoplasm, while at 25°C, proteins involved with cilium were upregulated in both strains. In La, the cytoskeleton was enriched, showing more regulation at Sta25, while in Lc, the peroxisome and lysosome were enriched at Sta34 and Sta25, respectively (**Supplementary Fig. 10C**).

**Supplementary Fig. 10.**
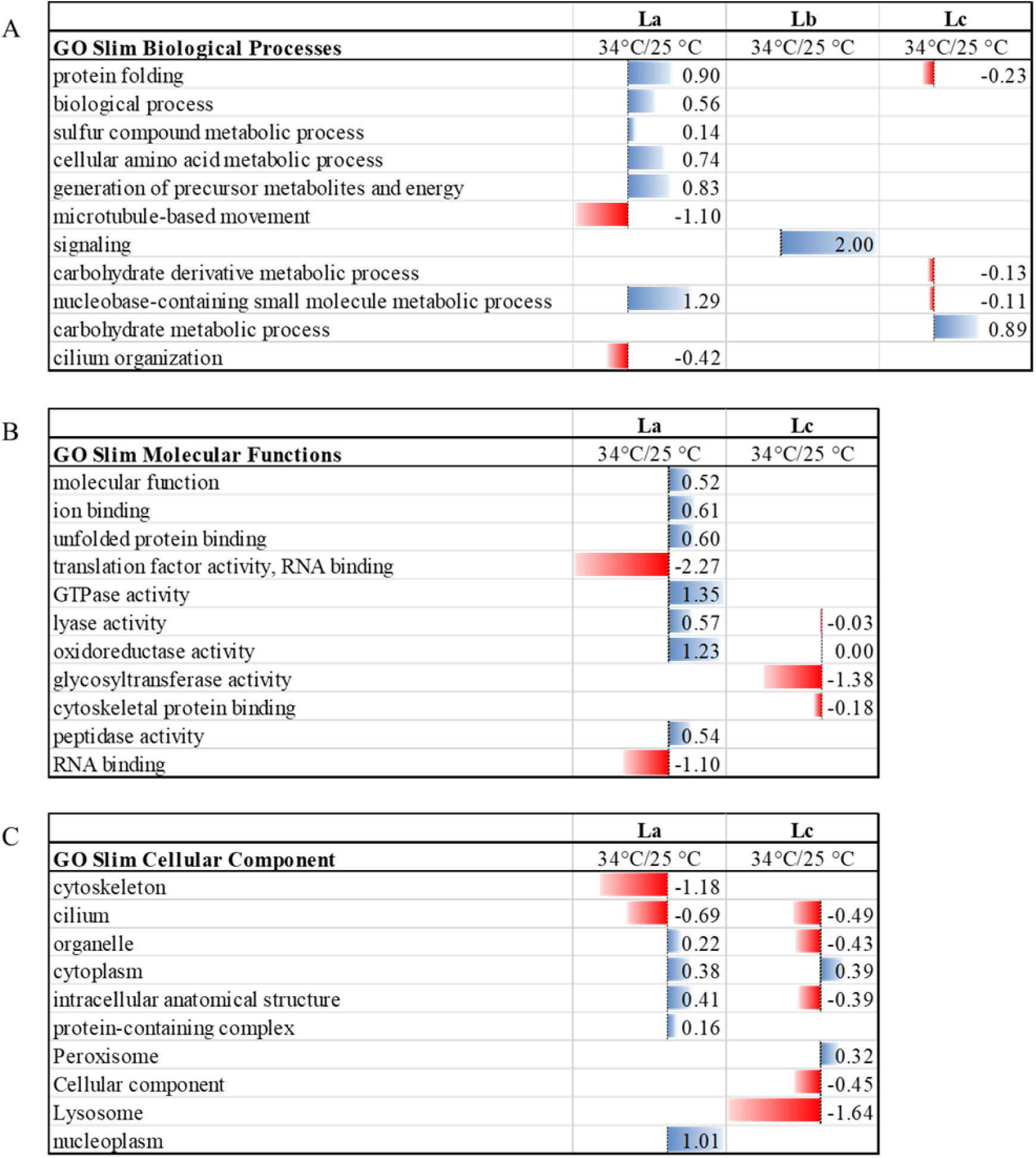
Gene ontology analysis combined with quantitative proteomics data of differentially expressed proteins between 25°C and 34°C for *L. amazonensis*, *L. braziliensis* and *L. infantum.* GO slim terms for Biological processes (**A**), Molecular functions (**B**), and Cellular compartments (C) are illustrated. Blue bars represent proteins upregulated in the stationary phase at 34°C, while red bars represent proteins upregulated in the stationary phase at 25°C.

In order to validate specific protein candidates identified by quantitative shotgun proteomics, a western blot assay was performed for α-tubulin, gp63, and HSP70 in the same species under the three conditions (**Fig. 6**). α-tubulin protein was identified in the three conditions of La and Lb. In both *Leishmania* species, increased expression of α-tubulin in the stationary phases compared to the exponential was shown (**Fig. 6A**). Moreover, temperature shift from 25°C to 34°C induced higher expression of α-tubulin.

**Fig. 6.**
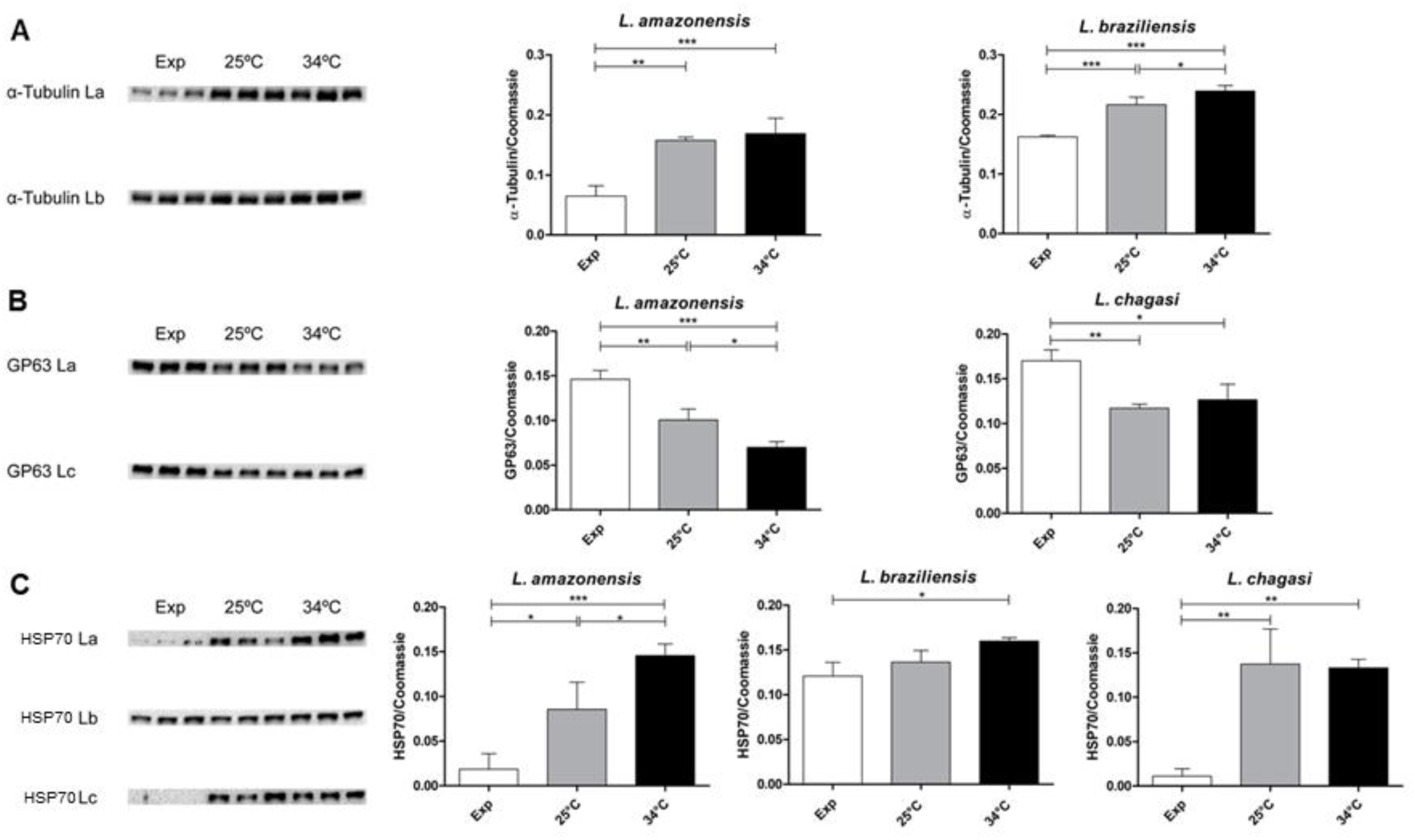
Differential expression of α-tubulin (A), GP63 (B) and HSP70 (C) in total protein extracts of promastigotes from exponential (Exp) phase, Sta25 (25°C), and Sta34 (34°C). Western blot was performed with biological triplicates for La = *L. amazonensis*, Lb = *L. braziliensis* and Lc = *L. infantum*. Data represented as mean±standard deviation. Values were considered significantly different for p-values<0.05 analyzed by one-way ANOVA.

Mass spectrometry-based proteomics analysis of α-tubulin for La revealed the differential expression of A4H727 and E9AP62 that was more expressed in the exponential compared to the stationary phase at 25°C and 34°C. No regulated α-tubulin protein was found for Lb or Lc (**Supplementary Fig. 6A).**

Western blot assay analysis of gp63 in the three conditions for each *Leishmania* species showed a decrease in the expression from exponential to stationary phases (**Fig. 6B**). In La, a thermal shift to 34°C showed a reduced gp63 expression compared to 25°C, while in Lc, the two values were not statistically different. For gp63, eight proteins were identified by mass spectrometry-based proteomics (**Supplementary Fig. 6B).** Q6LA77 (GP63-2), A4HUG0 (GP63-3), A4H638, E9AN54 and E9AN53 were detected in the three conditions in the three *Leishmania* species, while the other gp63 proteins are specific for either two *Leishmania* species in the different conditions. In La, Q6LA77, E9AN53, E9AN54, E9AN57 were upregulated in the exponential phase compared to Stat25 and Stat34, while A4HUF6 and E9AZL8 showed less expression at Exp compared to Sta25 and Sta34 conditions. This result suggests that the antibody used in the western blot has an epitope that recognizes an amino acid sequence similar to gp63 Q6LA77, E9AN53, E9AN54, E9AN57, which would explain the compatibility of the results. In Lb, no regulation was identified for the identified gp63 proteins. In Lc, A4HUG0, and E9AN54 were found regulated, being more expressed in the stationary phase (both Sta25 and Sta34) compared to the exponential phase, while E9AN53 was more expressed at Sta25 compared to Exp and Sta34 (25>Exp=34) (**Supplementary Fig. 6B).**

Western blot analysis of HSP70 in the three experimental conditions for each species revealed an increased expression from the exponential to the stationary phase (**Fig. 6C**). However, species-specific regulation levels were observed, with La and Lc showing the highest values between exponential and stationary phase at 25 and 34°C. Temperature shift from 25°C to 34°C of *Leishmania* parasites in the stationary phase significantly increased the expression of HSP70 in La (**Fig. 6C**). Mass spectrometry-based proteomic analysis allowed the identification of eight HSP70 proteins, of which seven were regulated (**Supplementary Fig. 6E-G).** In La, regulated HSP70 proteins A4HRI7 and A4H397 were least expressed in the exponential phase compared to stationary phases (25>34>Exp), E9AJF1 was more abundant at Sta25 but with similar expression at Exp and Sta34 (25>34=Exp), while A4I412 was more abundant in Exp and Sta34 phases compared to Sta25 (Exp>34>25). No regulation of HSP70 proteins was identified in Lb. In Lc, six HSP70 proteins were regulated, with E9B099 and A4H397 having less expression at the exponential phase compared to the two stationary phases (34=25>Exp). On the other hand, A4I412, A4HGY1 and A4I253 were more expressed in the exponential phase when compared to the two stationary conditions (Exp>34=25), while E9AJF1 was more abundant in Sta34 and least in Sta25 (34>Exp>25). These data demonstrate a species-specific HSP70 profile of the three *Leishmania* species (**Supplementary Fig. 6E**). These data confirmed the upregulation of more HSP70 proteins in the stationary phases compared to the exponential phase.

Analysis of the parasites’ metabolic function corresponding to cell viability was measured by MTT colorimetric assay, revealing a higher activity at 25°C compared to 34°C (**Fig. 7A**). Upon thermal stress, Lb had pronounced decrease in cell viability/metabolic activity compared to La and Lc, suggesting a species-specific differential response to heat stress (**Fig. 7A**). This result is in apparent contrast to the negligible proteome modulation in Lb from 25°C to 34°C (**Fig.5B, Supplementary Fig. 8D**), which showed the regulation of only nine proteins, compared to 1380 and 651 proteins in *L. amazonensis* and *L. infantum*, respectively, calling attention to the possible regulation at post-transcriptional level or modulation of the proteome at the post-translation level upon heat stress in *L. braziliensis*. Proteome modulation at the PTM level, such as phosphorylation, acetylation, methylation and glycosylation, which play key roles in stage-specific signal transduction pathways, has been described in *Leishmania* species during stage transition and temperature changes [50-54]. Lc species showed more resistance to thermal stress than La and Lb. This result can be associated with Lc being one of the species that causes visceral leishmaniasis. The expression of heat shock proteins was evaluated in La, Lb and Lc under the three different conditions. Of the 40 heat shock proteins identified, 22 were regulated in La, Lb and Lc with similarities and differences, as highlighted in the HSP expression profiles (**Fig. 7B, C, D**). In La, Exp and Sta34 phases clustered close together, well separated from Sta25 based on regulated HSP expression profiles, where 9 HSPs were highly expressed in the Exp and Sta34 phases and 7 HSPs were exclusively upregulated in Sta25 (**Fig. 7B**). HSPs upregulated in Exp and Sta34 included HSP60, mitocnhondrial HSP60, HSP70, while HSPs upregulated in Sta25 included HSP60, mitochondrial HSP60, HSP70, HSP78, HSP83-1, and HSP100 (**Fig. 7B**). One HSP83, A4HL69, was exclusively missing in Sta25, while another HSP83-1, E9AHM9, was upregulated in Sta34 compared to Sta25 and Exp (34>25>Exp). Regulated HSPs in Lb showed a clustering different from La, with Exp being separated from the two stationary phases (**Fig. 7C**). Ten (10) HSPs, including HSP60, mitochondrial HSP60, HSP70, and HSP 83 were upregulated at the exponential phase compared to Sta25 and Sta34, while one mitochondrial HSP60, E9ASX7 was found exclusively expressed at the exponential phase. The separation between the biological triplicates at the stationary phases was not evident, which could have been influenced by the NaN values for several HSPs in Lb. Contrary, regulated HSPs in Lc showed a clear separation between the two stationary phases, which were separated from the exponential phase (**Fig. 7D**). Only this clustering was similar to that observed for the total regulated proteome (**Fig. 5C**). In Lc, 10 HSPs were upregulated in Sta34 compared to Sta25, including HSP70, mitochondrial HSP60, HSP83-1 and HSP100.

**Fig. 7.**
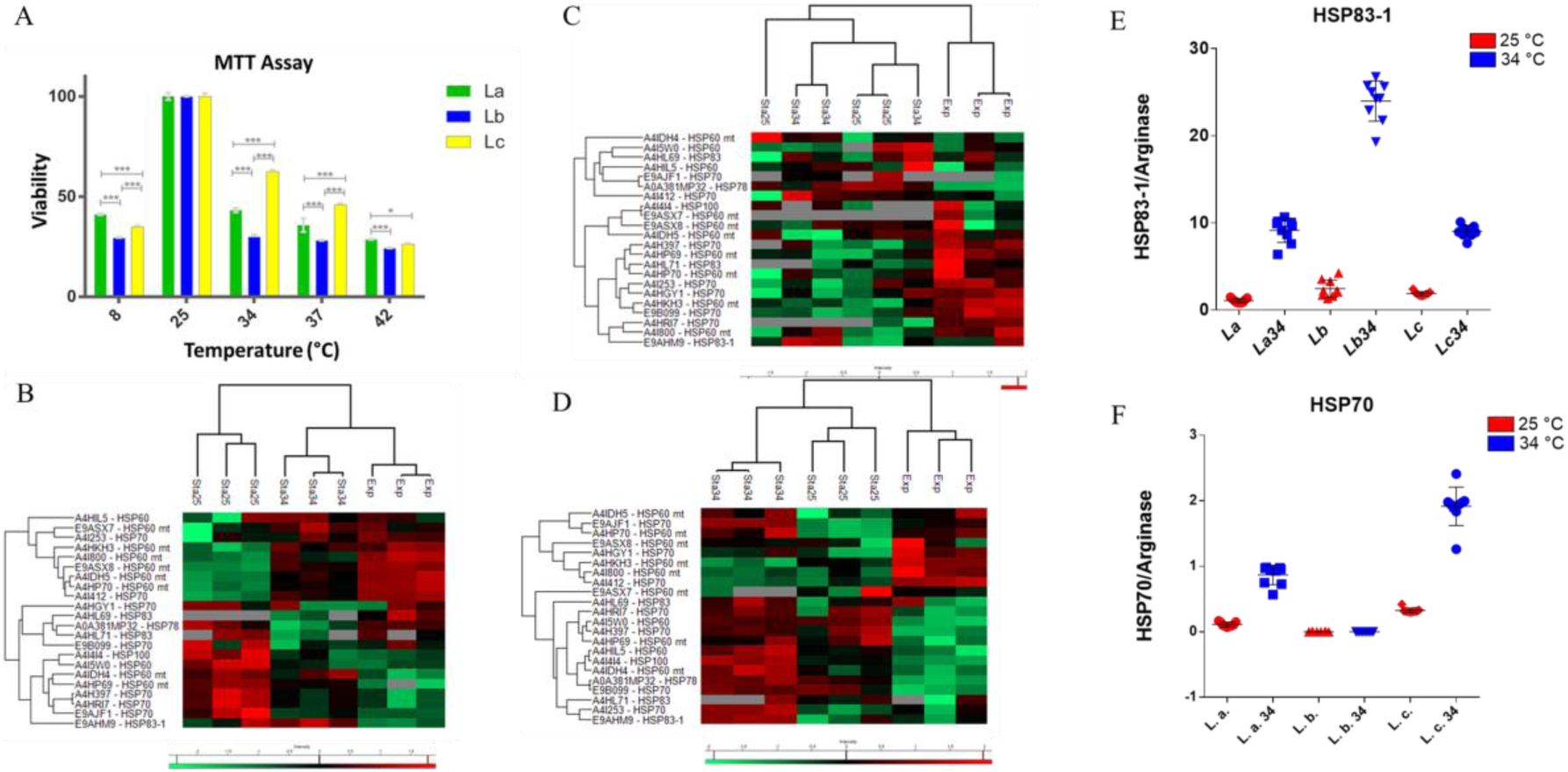
*Leishmania* parasite viability and HSP expression profiles at different temperatures. (**A**) Viability of promastigotes in the stationary phase of La = *L. amazonensis*, Lb = *L. braziliensis*, and Lc = *L. infantum* after 24 h at different temperatures. The viability analysis was done by MTT assay, and ANOVA statistical test was performed (* p <0.05; *** p <0.001). The graph is representative of five independent experiments. Heatmaps of the Heat shock proteins (HSPs) identified and quantified by nLC-MS/MS, and their regulation in the species La (**B**), Lb (**C**) and Lc (**D**) in the three conditions. The red color represents the most expressed proteins for each species, the least expressed by the green color and absent in gray. RT-qPCR was used to evaluate the transcript abundances for (**E**) HSP 83-1 and (**F**) HSP70.

Analysis by RT-qPCR showed that HSP83-1 mRNA expression was increased in the three species upon temperature shift being more regulated in Lb (**Fig. 7D**). Moreover, HSP70 mRNA expression was increased in the La and Lc, being more in Lc, but was not regulated in Lb (**Fig. 7E**).

## DISCUSSION

Proteomic approaches have improved the current understanding of *Leishmania* biology. In particular, protein-based molecular changes at different stages of *Leishmania* development, *Leishmania*-host interaction, drug resistance, disease pathogenesis, and potential biomarkers have been elucidated by proteomics [10, 55]. In this study, we analyzed the proteome modulation of three New World *Leishmania* species belonging to the subgenera *Leishmania* (*L. amazonensis* and *L. infantum-syn L. chagasi*) and *Viannia* (*L. braziliensis*) [56].

Among the *Leishmania* subgenus, *L. amazonensis* and *L. infantum* belong to the *L. (Leishmania) mexicana* and *L. (Leishmania) donovani* complexes, respectively. The disease outcome in human hosts differs for the three species, where *L. amazonensis* and *L. braziliensis* are dermotropic, usually causing localized cutaneous lesions that may develop into diffuse and mucocutaneous, respectively, while *L. infantum* is viscerotropic, causing visceral leishmaniasis. Genetic variability, phenotypic and clinical manifestation associated with *L. braziliensis* have been demonstrated in several strains isolated in different geographical regions [57, 58]. The clinical outcome of leishmaniasis is driven by the difference between species in conjunction with host genetics and their immune response. Although the information on host immune response against different *Leishmania* species has made great strides, the factors contributing to these differences are largely unknown [59, 60], despite several differences that have been reported among species. Indeed, amastigotes from *Leishmania* species belonging to the *L. mexicana* complex, such as *L. amazonensis*, are housed in large numbers within a large parasitophorous vacuole within the macrophages [61, 62]. In contrast, amastigotes from *L. donovani* and *L. braziliensis* complexes reside intracellularly within small parasitophorous vacuoles that typically contain a single parasite [63, 64]. *Leishmania* from *Viannia* and *Leishmania* have important differences in their development in the vector. They also have differences in RNAi machinery and a higher divergence in their genomes. In the phlebotomine, the differentiation of *Leishmania* from procyclic promastigotes to metacyclic promastigotes prepares the parasite for transmission to a mammalian host. Molecular characteristics (genes, proteins, and metabolites) among three *Leishmania* species have been investigated to understand the causes of their clinical peculiarities. This study implemented a comprehensive experimental approach based on large-scale quantitative proteomics and orthogonal functional assays. Cell viability of the three species under the three conditions was evaluated and provided the basis for interpreting proteomic differences due to the remodeling of intracellular signaling pathways rather than cell death. Specific modulation of protein classes of selected *Leishmania* species during growth (exponential phase, and stationary phase), and temperature changes (change from 25°C to 34°C) was observed. The first objective of this study was to compare three *Leishmania* species to identify differences in their proteome correlated with their pathophysiological outcome. A total of 7047 proteins were identified, considering the three species and the three growing conditions. To our knowledge, this number of identified proteins represents the most extensive data set reported to date. Analyzing the total number of proteins identified for each *Leishmania* species under the three conditions, *L. braziliensis* presented the highest number of proteins, followed by *L. infantum* and *L. amazonensis*. The highest number of proteins identified for the three species was in the exponential phase, followed by the stationary phase at 34°C, and the stationary phase at 25°C. The lower number of peptides and proteins for the species *L. amazonensis* in all conditions may have been caused by using the *L. mexicana* database available from the UniProt public database. Unfortunately, a protein database for *L. amazonensis* was not found in UNIPROT, and although the TriTrypDB database contains a list of *L. amazonensis* proteins, these data are from ESTs and ORFs and are not well noted, according to the criteria used in this study.

About 24173 proteins from the *Leishmania* species were used for the analysis, and the same database was used to search the mass spectrometry data for the three species, under the three conditions. The three species mentioned above were selected to represent the complex of each species studied. Due to the high similarity of protein sequences among the three species, some protein groups were identified based on shared peptides.

One of the first reports using proteomic approaches to the *Leishmania* study explored 2D electrophoresis (2DE) of New World *Leishmania* promastigotes in the exponential phase belonging to the species complex *L. mexicana* and *L. braziliensis* [65]. A different protein pattern was observed for the two complexes with minor differences within the same complex, indicating a species-specific proteome map [65]. The proteomic comparison of *L. tropica*, *L. major*, and *L. infantum* promastigotes was performed using 2DE combined with LC-MS/MS, and the raw data were searched against the *L. major* protein database [66]. Regulated proteins were involved in the cytoskeleton, oxidative stress, proteolysis, and transport. Cluster analysis based on protein differential expression showed that the two clustered cutaneous species separated from the visceral, suggesting a modulation of the proteome specific for the disease outcome, although the promastigote stage has been analyzed [66]. Another study analyzed the membrane proteome of *L. mexicana* and *L. infantum* in the promastigote and amastigote stages. Within the regulated proteins, gp63 was upregulated at the promastigote stage in both species, while kinetoplastid membrane protein-11 was upregulated at the promastigote stage in *L. infantum*, being less regulated at *L. mexicana* [67]. These data show a specific modulation of the species proteome during parasite development. Analyzing the percentage of regulated proteins, we detected up to 40% of regulated proteins in *L. amazonensis*. In other studies, the transition from promastigote to amastigote revealed that 5-10% of the proteome is differentially regulated [34, 68-70]. The data presented here show a higher regulation from exponential to stationary growth phases.

Previous proteomic studies of *Leishmania* spp. in transition from different stages of life have been reported. A quantitative protein analysis over time in *L. donovani* during axenic differentiation from promastigote to amastigote revealed lower expression of anabolic functions, such as protein translation, and higher expression of catabolic processes TCA cycle and β-oxidation [71]. Under axenic conditions, promastigotes undergo drastic changes in pH (from 7 to 5) and temperature (from 25°C to 37°C). These conditions reproduce some of the characteristics of the lysosomal environment of amastigotes residing within macrophages. Interestingly, in the first 10 hours of promastigote exposure, few proteins changed significantly, indicating a gradual and regulated remodeling of intracellular pathways at the protein level [71]. Analysis of the transcripts revealed early modulation compared to protein level under the same experimental conditions [12]. The present study investigated temperature shifts from 25°C to 34°C. Temperature shift is important for isolating the effect of temperature on *Leishmania* proteome modulation, as discussed below.

Another study using mass spectrometry to identify *Leishmania* proteins identified 3700 proteins in *L. major* [72]. Studies with *L. donovani* using 2DE gel and MALDI-TOF were also performed. In one, the soluble proteins were analyzed to identify potential antigens [73], while the other study identified 132 proteins from clinical isolates [74]. There are also studies to identify *Leishmania* phosphoproteome. Two studies using *L. donovani* axenic promastigotes and amastigotes identified 171 proteins [52], and 282 proteins [75]. Our study expands the repertoire of proteins identified and quantified in the species *L. amazonensis*, *L. braziliensis*, and *L. infantum* offering an important database for the scientific community.

The second objective is related to understanding the differences between exponential and stationary promastigotes in three *Leishmania* species fostering our knowledge of the molecular changes in the developmental stages in the phlebotomine sand fly vector. To our knowledge, there are no published studies on the quantitative mass-spectrometry-based proteomic analysis of *Leishmania* spp., comparing the exponential and stationary growth phases. However, a 2DE coupled to MALDI-TOF proteomic approach was applied to the early stages of exponential and stationary phases of *L. infantum*, showing 28 proteins differentially regulated between the two stages [76]. In addition, the proteome profiles of *L. infantum* exosomes at the exponential and stationary growth phases were recently described, showing stage-specific exosome protein profiles, and a notably increased level of proteins in the exponential phase [77]. Recently, a deep quantitative proteomic analysis of *Leishmania Viannia* species, such as *L. braziliensis*, *L. panamensis*, and *L. guyanensis,* was performed, elucidating the copy numbers of 7000 proteins [78]. Species-specific regulation of metabolic pathways was demonstrated. Moreover, metabolomic studies in *L. donovani* and *L. major* have been reported. Untargeted metabolomic analysis of the transition from the exponential to the stationary phase revealed changes in specific lipid classes such as glycerophospholipids, sphingolipids, and glycerolipids. ^1^H NMR-based metabolomic analysis during the transition from the exponential to the stationary phase induced changes in specific metabolites such as citraconic acid, isopropyl acid, L-leucine, ornithine, caprylic acid, capric acid, and acetic acid [79, 80]. Moreover, a modification of the structure of lipophosphoglycan during *L. major* differentiation from the procyclic promastigote to the metacyclic promastigote stage was described [15].

The surface modulation of the proteome during the transition from exponential to stationary phase in *L. infantum* showed increased resistance to complement-mediated lysis. Exponential phase promastigotes from *Leishmania* parasites modify morphological characteristics after incubation with normal human serum, inducing the deposition of C3, C5, and C9 proteins on the parasite cell membrane [81]. In our study, the three New World *Leishmania* species showed protein regulation during the exponential to stationary transition. In particular, *L. infantum* presented the highest percentage of regulated proteins (40.3%), followed by *L. amazonensis* (40%) and *L. braziliensis* (21.5%). During the exponential to stationary transition, species-specific proteome regulation indicates that *L. infantum* and *L. amazonensis* are more affected than *L. braziliensis* during their insect-vector development.

The third objective was focused on understanding the molecular plasticity of the three species submitted to thermal stress. In order to address the effect of temperature on parasites, stationary phase promastigotes grown at 25°C were incubated at 34°C for 24 hours. Thermal stress induced changes in the parasite morphology. The parasite’s body becomes rounded and a smaller flagellum is observed on higher temperature incubation. Temperature is one of the triggering factors for parasite modifications, allowing it to adapt to the mammalian host. The parasite’s thermal shock response helps in maintaining cellular homeostasis and survival, and heat shock proteins are a crucial player in this response [82]. Proteomic analysis of the three conditions for each species showed a higher number of proteins for the exponential phase, followed by the stationary phase at 34°C and least in the stationary phase at 25°C. The decrease in the number of proteins between the exponential and stationary phase at 25°C is related to the transition from proliferative to quiescent growth stages.

Interestingly, *Leishmania* parasites collected on the 6th day of cultivation and incubated for 24 hours at 34°C increased the number of proteins detected, showing modulation of intracellular proteome dictated by temperature shift. *L. amazonensis* presented the highest number of regulated proteins compared to *L. infantum* between the stationary phases at 25°C and 34°C, while *L. braziliensis* showed minimal regulation upon thermal stress. This result shows, again, the species-specific response to heat stress. Gene ontology analysis showed the abundance of proteins with enrichment in unfolded protein binding in *L. amazonensis* followed by *L. braziliensis* compared to *L. infantum* during temperature shift from 25°C to 34°C (La > Lb > Lc), suggesting higher levels of protein misfolding in *L. amazonensis* and *L. braziliensis* compared to *L. infantum* at higher temperatures. During temperature shift, GO analysis showed higher enrichment of RNA binding proteins in *L. amazonensis* and *L. infantum* compared to *L. braziliensis,* suggesting species-specific and temperature-dependent modulation of gene expression. Using orthogonal approaches, all comparisons were further validated by focusing on α-tubulin, gp63, and HSPs. These protein families were chosen due to their importance in parasite morphology and virulence. Trypanosomatids’ cytoskeleton is responsible for maintaining the parasite’s shape throughout its life cycle, cell division, and intracellular charge transport [83]. The cytoskeleton, an interconnected network of filamentous polymers and regulatory proteins, is composed of microtubules, actin filaments, and intermediate filaments. Microtubules, the largest structure and major component of the cytoskeleton are comprised of α and β-tubulin subunits assembled on linear protofilaments [84]. Parasites are exposed to various osmotic changes, from extreme fluctuations in the osmotic force of the vector’s midgut to acidic phagolysosomes, and the reorganization of the cytoskeleton allows them to control their shape. In this study, we investigated the expression of α-tubulin protein among the different *Leishmania* species under growth conditions and temperature changes. Western blot analysis showed a higher expression of α-tubulin in the stationary phase at 34°C compared to the other growth conditions and in *L. braziliensis* compared to the other species. These results agree with mass spectrometry data that reported α-tubulin (AH4727) with the highest expression in *L. braziliensis*.

The genomic organization of α-tubulin genes in *L. braziliensis* is different compared to *L. major* and *L. infantum*, with a common characteristic of several gene copies arranged together [85]. The α-tubulin gene in *L. major* is located in a locus on chromosome 13 with 12 copies with the identical ORFs arranged in series [86]. The α-tubulin gene in *L. infantum* exists in chromosome 13, one with a single copy and the other with a complete and truncated copy. The α-tubulin gene in *L. braziliensis* exists at two *loci* on chromosomes 13 and 29. The locus on chromosome 13 contains two complete copies and the α-tubulin-like gene. The locus on chromosome 29 contains a copy with a truncated form. Regarding *Leishmania* stages, the tubulin mRNA level was higher in promastigotes than in amastigote forms. This result was also observed in *L. enrietti* and *L. major* [87, 88]. However, no difference was observed in *L. mexicana* between promastigote and amastigote stages [89]. In addition, temperature-dependent regulation was observed for α-tubulin mRNA. *L. braziliensis* promastigotes incubated at 35°C showed decreased α-tubulin mRNA expression [85]. This trend has been demonstrated at the protein level with decreased tubulin expression during the transition from promastigote to amastigote forms [89-91]. The data obtained in this study show that the regulation of α-tubulin is dependent on *Leishmania* species and temperature.

*Leishmania* proteases play an important role in various biological processes such as cytoadherence, virulence, invasion and spread to host tissues, survival in different host environments, and immune system inhibition [92-94]. Cysteine, serine, aspartic acid, and metalloproteases were mapped in the *Leishmania* genome and identified at the protein level [95, 96]. Gp63, also known as leishmanolysin, or promastigote surface protease, is the most abundant surface protease in many *Leishmania* spp [97, 98]. It is a zinc-dependent metalloprotease with broad substrate specificity cleaving multiple intracellular proteins, modifying the host nuclear envelope, and degrading extracellular matrix proteins [27, 99]. In addition, gp63 has important functions as a virulence factor and complement fixation [25, 26]. Gp63 is expressed at higher levels in promastigote and at lower levels in amastigote life stages [24]. However, conflicting results have been obtained using antibodies against different epitopes of the molecule [100, 101]. Multiple copies of *Leishmania* gp63 genes are reported together at a single chromosome site [102, 103] having a large polymorphism within the same species [104], which may explain the difference between the results obtained in this study through proteomics and Western blot analysis. Inclusion of protease inhibitors in the lysis buffer did not affect the activity of gp63, corroborating gp63’s resistance to majority of protease inhibitors previously reported [97, 105], and the analysis of the proteolytic activity of the three *Leishmania* species under the three conditions revealed a specific protease pattern for each species and condition. Early reports of the proteolytic profile of five *L. braziliensis* strains isolated from patients in Colombia and Brazil with different clinical outcomes revealed 50-125 kDa proteins [106]. Promastigote extracts and their supernatants were tested against fish gelatin. These proteases were inhibited by 1,10-phenanthroline, indicating a metalloprotease activity [106]. *L. braziliensis* pattern showed four major bands between 60 and 80 kDa. The lower molecular weight band at 63kDa of *L. braziliensis* was found to increase at the stationary phase at 34°C. These results confirm the differential expression and activity of gp63 in *Leishmania* spp., under different conditions. Along with α-tubulin, the gp63 family regulation is another important molecule critical for investigation as a possible therapeutic target.

Heat shock proteins (HSPs) showed differential expression between different *Leishmania* species and growing conditions. HSPs are highly conserved proteins constitutively transcribed in all cell types and are regulated at the post-transcriptional level [107]. Their primary structures and molecular mass divide them into different families such as HSP110, HSP104/100, HSP90/83, HSP70, HSP60, HSP40, HSP10, and small heat shock proteins. They have the function of interacting with various proteins controlling their maturation, activation, translocation, and degradation [32, 108]. HSP family 70 (HSP70) is numerous and diverse in *Leishmania* spp., [32]. HSP70, HSP90 and HSP40 are known to be part of the complex involved in the correct folding, maturation and assembly of peptides [109]. Heat shock protein family 90, also known as HSP83, plays an important role in controlling parasite life. HSP83 is an ATP-dependent molecular chaperone. This protein is not involved in the protection of the parasite against environmental stress. Indeed, although there is an increase in its expression in thermal stresses, it is also abundant under normal growth conditions, as in the proliferation of the parasite within the phlebotomine sand flies (promastigote) or within mammalian hosts (amastigote). The transcriptome levels in *L. major* promastigotes are modulated during heat shock treatment for 2h at 37°C, with 18% and 20% of the transcripts being up and downregulated, respectively [110]. This study showed the upregulation (fold change > 2) of transcripts coding for HSP70-I, two HSP83/90, HSP100, and two small HSPs at 37° compared to 26°C. Moreover, upregulated mRNA transcripts coding for proteins involved in such GO processes as calcium ion homeostasis, modulation by symbiont of host phagocytosis, active evasion of host immune response via regulation of host complement system and the modulation of host nitric oxide-mediated signaling pathways, while the downregulated transcripts code for proteins involved in ribosome biogenesis and translation [110]. In our study, several HSPs were found regulated between the different stages of *Leishmania* and thermal stress. In particular, HSP70 showed higher expression in the exponential phase using mass spectrometry-based proteomics. Western blot analysis showed higher expression in the stationary phase and increased even more with the temperature change. This specific regulation of HSPs belonging to different families, detected by quantitative proteomics, opens new opportunities to study their influence as specific potential therapeutic targets.

It is important to mention that some discrepancies between protein expression (by western blot) and proteomic data were observed, especially for α-tubulin protein (**Fig. 6A** and **Supplementary Fig. 6A**). Using antibody-based assays to quantify proteins in different conditions is a well-established technique that has been applied worldwide to different organisms. One of the biggest challenges in western blot analysis is antibody specificity. This issue becomes important for analyzing proteins belonging to multi-gene families with subtle sequence differences. Furthermore, comparing species of the same organism represents another challenge related to species-specific differences in protein sequence based on phylogenetic relationships. For *Leishmania*, the number and quality of antibodies available are scarce compared to humans or mice. In most cases, these antibodies are poorly characterized, and the knowledge of their antigen-epitope is private and unavailable. In addition, a reference sample for antibody specificity testing is not available in most cases. The bottom-up mass spectrometry approach, used in this manuscript, allows simultaneous monitoring of thousands of reliably identified and quantified proteins. This method allows high confidence in identifying and quantifying proteins belonging to multigene families.

## CONCLUSION

The quantitative proteome of *Leishmania (Leishmania) amazonensis, Leishmania (Viannia) braziliensis*, and *Leishmania (Leishmania) infantum* in this study highlight the differences and similarities among three species that may be linked to their phylogenetic proximity, phenotypes, and pathological outcome. The proteomic differences identified for the growth phase transition and heat stress consolidate the concept of proteomic plasticity of the three species, reshaped over external stimulus induced by selective host pressure on the parasite. Many of the proteins identified in this study are associated with biological processes essential for parasite survival and infectivity. Among these proteins, α-tubulin, gp63 and HSPs showed a specific pattern that deserves further investigation. Different proteins belonging to each protein family presented a specific expression pattern in the analyzed species and conditions. Following bibliographic research on proteomic studies on *Leishmania* parasites under different conditions, it is clear that α-tubulin, gp63, and HSPs are among the most commonly identified proteins that have been proposed as drug targets and vaccine candidates. The constant identification of these proteins may be associated with their abundance within *Leishmania* parasites and their essential functions.

## Acknowledgments

The work was supported by grants and fellowships from FAPESP (2018/18257-1, 2018/15549-1, 2020/04923-0 to GP; 2017/04032-5 and 2021/14751-4 to SNM; 2016/23689-2 to JSS; 2018/13283-4 to GSDO; 2021/08915-4 to BSS and 2020/13562-0 to MC), from *Coordenação de Aperfeiçoamento de Pessoal de Nível Superior* – Brasil (CAPES) (Código de Financiamento 001 to LR-F; to DOQ; to IPS and; to DBA) and from *Conselho Nacional de Desenvolvimento Científico Tecnológico* (CNPq) (“bolsa de produtividade” to GP).

## References

[1] G. Grimaldi, Jr., R.B. Tesh, Leishmaniases of the New World: current concepts and implications for future research, Clinical microbiology reviews 6(3) (1993) 230–50.

[2] L.C. Castellucci, L.F. Almeida, S.E. Jamieson, M. Fakiola, E.M. Carvalho, J.M. Blackwell, Host genetic factors in American cutaneous leishmaniasis: a critical appraisal of studies conducted in an endemic area of Brazil, Memorias do Instituto Oswaldo Cruz 109(3) (2014) 279–88.

[3] O.E. Akilov, A. Khachemoune, T. Hasan, Clinical manifestations and classification of Old World cutaneous leishmaniasis, International journal of dermatology 46(2) (2007) 132–42.

[4] H. Goto, J.A. Lauletta Lindoso, Cutaneous and mucocutaneous leishmaniasis, Infectious disease clinics of North America 26(2) (2012) 293–307.

[5] B.S. McGwire, A.R. Satoskar, Leishmaniasis: clinical syndromes and treatment, QJM: monthly journal of the Association of Physicians 107(1) (2014) 7–14.

[6] Y. Hashiguchi, E.L. Gomez, H. Kato, L.R. Martini, L.N. Velez, H. Uezato, Diffuse and disseminated cutaneous leishmaniasis: clinical cases experienced in Ecuador and a brief review, Trop Med Health 44 (2016) 2.

[7] S.T. de Macedo-Silva, J.A. Urbina, W. de Souza, J.C. Rodrigues, In vitro activity of the antifungal azoles itraconazole and posaconazole against Leishmania amazonensis, PloS one 8(12) (2013) e83247.

[8] J. Alvar, I.D. Vélez, C. Bern, M. Herrero, P. Desjeux, J. Cano, J. Jannin, M. den Boer, Leishmaniasis worldwide and global estimates of its incidence, PloS one 7(5) (2012) e35671.

[9] P.A. Bates, Revising Leishmania’s life cycle, Nature microbiology 3(5) (2018) 529–530.

[10] J. Capelli-Peixoto, S.N. Mule, F.T. Tano, G. Palmisano, B.S. Stolf, Proteomics and Leishmaniasis: Potential Clinical Applications, Proteomics. Clinical applications 13(6) (2019) e1800136.

[11] L.M. De Pablos, T.R. Ferreira, P.B. Walrad, Developmental differentiation in Leishmania lifecycle progression: post-transcriptional control conducts the orchestra, Current opinion in microbiology 34 (2016) 82–89.

[12] T. Lahav, D. Sivam, H. Volpin, M. Ronen, P. Tsigankov, A. Green, N. Holland, M. Kuzyk, C. Borchers, D. Zilberstein, P.J. Myler, Multiple levels of gene regulation mediate differentiation of the intracellular pathogen Leishmania, FASEB journal: official publication of the Federation of American Societies for Experimental Biology 25(2) (2011) 515–25.

[13] S.J. Turco, S.R. Hull, P.A. Orlandi, Jr., S.D. Shepherd, S.W. Homans, R.A. Dwek, T.W. Rademacher, Structure of the major carbohydrate fragment of the Leishmania donovani lipophosphoglycan, Biochemistry 26(19) (1987) 6233–8.

[14] M.J. McConville, A. Bacic, G.F. Mitchell, E. Handman, Lipophosphoglycan of Leishmania major that vaccinates against cutaneous leishmaniasis contains an alkylglycerophosphoinositol lipid anchor, Proceedings of the National Academy of Sciences of the United States of America 84(24) (1987) 8941–5.

[15] M.J. McConville, S.J. Turco, M.A. Ferguson, D.L. Sacks, Developmental modification of lipophosphoglycan during the differentiation of Leishmania major promastigotes to an infectious stage, The EMBO journal 11(10) (1992) 3593–600.

[16] E.M. Saraiva, P.F. Pimenta, T.N. Brodin, E. Rowton, G.B. Modi, D.L. Sacks, Changes in lipophosphoglycan and gene expression associated with the development of Leishmania major in Phlebotomus papatasi, Parasitology 111 (Pt 3) (1995) 275–87.

[17] R.P. Da Silva, B.F. Hall, K.A. Joiner, D.L. Sacks, CR1, the C3b receptor, mediates binding of infective Leishmania major metacyclic promastigotes to human macrophages, Journal of immunology (Baltimore, Md.: 1950) 143(2) (1989) 617-22.

[18] D.M. Mosser, A. Brittingham, Leishmania, macrophages and complement: a tale of subversion and exploitation, Parasitology 115 Suppl (1997) S9–23.

[19] E.D. Franke, P.B. McGreevy, S.P. Katz, D.L. Sacks, Growth cycle-dependent generation of complement-resistant Leishmania promastigotes, Journal of immunology (Baltimore, Md.: 1950) 134(4) (1985) 2713-8.

[20] S.M. Puentes, R.P. Da Silva, D.L. Sacks, C.H. Hammer, K.A. Joiner, Serum resistance of metacyclic stage Leishmania major promastigotes is due to release of C5b-9, Journal of immunology (Baltimore, Md.: 1950) 145(12) (1990) 4311-6.

[21] P.F. Pimenta, E.M. Saraiva, D.L. Sacks, The comparative fine structure and surface glycoconjugate expression of three life stages of Leishmania major, Experimental parasitology 72(2) (1991) 191–204.

[22] S.J. Turco, D.L. Sacks, Expression of a stage-specific lipophosphoglycan in Leishmania major amastigotes, Molecular and biochemical parasitology 45(1) (1991) 91–9.

[23] S.N. Mule, J.S. Saad, L.R. Fernandes, B.S. Stolf, M. Cortez, G. Palmisano, Protein glycosylation in Leishmania spp, 16(5) (2020) 407–424.

[24] T.O. Frommel, L.L. Button, Y. Fujikura, W.R. McMaster, The major surface glycoprotein (GP63) is present in both life stages of Leishmania, Molecular and biochemical parasitology 38(1) (1990) 25–32.

[25] M. Olivier, V.D. Atayde, A. Isnard, K. Hassani, M.T. Shio, Leishmania virulence factors: focus on the metalloprotease GP63, Microbes and infection 14(15) (2012) 1377–89.

[26] A. Brittingham, C.J. Morrison, W.R. McMaster, B.S. McGwire, K.P. Chang, D.M. Mosser, Role of the Leishmania surface protease gp63 in complement fixation, cell adhesion, and resistance to complement-mediated lysis, Journal of immunology (Baltimore, Md.: 1950) 155(6) (1995) 3102-11.

[27] B.S. McGwire, K.P. Chang, D.M. Engman, Migration through the extracellular matrix by the parasitic protozoan Leishmania is enhanced by surface metalloprotease gp63, Infection and immunity 71(2) (2003) 1008–10.

[28] D. Zilberstein, M. Shapira, The role of pH and temperature in the development of Leishmania parasites, Annual review of microbiology 48 (1994) 449–70.

[29] H.L. Callahan, I.F. Portal, S.J. Bensinger, M. Grogl, Leishmania spp: temperature sensitivity of promastigotes in vitro as a model for tropism in vivo, Experimental parasitology 84(3) (1996) 400–9.

[30] K.W. Hunter, C.L. Cook, E.G. Hayunga, Leishmanial differentiation in vitro: induction of heat shock proteins, Biochemical and biophysical research communications 125(2) (1984) 755–60.

[31] M. Argaman, R. Aly, M. Shapira, Expression of heat shock protein 83 in Leishmania is regulated post-transcriptionally, Molecular and biochemical parasitology 64(1) (1994) 95–110.

[32] A. Hombach, G. Ommen, A. MacDonald, J. Clos, A small heat shock protein is essential for thermotolerance and intracellular survival of Leishmania donovani, Journal of cell science 127(Pt 21) (2014) 4762–73.

[33] J.B. de Jesus, C. Mesquita-Rodrigues, P. Cuervo, Proteomics advances in the study of Leishmania parasites and leishmaniasis, Sub-cellular biochemistry 74 (2014) 323–49.

[34] Y. El Fakhry, M. Ouellette, B. Papadopoulou, A proteomic approach to identify developmentally regulated proteins in Leishmania infantum, Proteomics 2(8) (2002) 1007–17.

[35] J. Walker, N. Acestor, R. Gongora, M. Quadroni, I. Segura, N. Fasel, N.G. Saravia, Comparative protein profiling identifies elongation factor-1beta and tryparedoxin peroxidase as factors associated with metastasis in Leishmania guyanensis, Molecular and biochemical parasitology 145(2) (2006) 254–64.

[36] C. Cassagne, F. Pratlong, F. Jeddi, R. Benikhlef, K. Aoun, A.C. Normand, F. Faraut, P. Bastien, R. Piarroux, Identification of Leishmania at the species level with matrix-assisted laser desorption ionization time-of-flight mass spectrometry, Clinical microbiology and infection: the official publication of the European Society of Clinical Microbiology and Infectious Diseases 20(6) (2014) 551–7.

[37] O. Mouri, G. Morizot, G. Van der Auwera, C. Ravel, M. Passet, N. Chartrel, I. Joly, M. Thellier, S. Jauréguiberry, E. Caumes, D. Mazier, C. Marinach-Patrice, P. Buffet, Easy identification of leishmania species by mass spectrometry, PLoS neglected tropical diseases 8(6) (2014) e2841.

[38] S. Sundar, B. Singh, Understanding Leishmania parasites through proteomics and implications for the clinic, Expert Rev Proteomics 15(5) (2018) 371–390.

[39] A. Parthasarathy, K. Kalesh, Defeating the trypanosomatid trio: proteomics of the protozoan parasites causing neglected tropical diseases, 11(6) (2020) 625–645.

[40] I.P. Sauter, K.G. Madrid, J.B. de Assis, A. Sá-Nunes, A.C. Torrecilhas, D.I. Staquicini, R. Pasqualini, W. Arap, M. Cortez, TLR9/MyD88/TRIF signaling activates host immune inhibitory CD200 in Leishmania infection, JCI Insight 4(10) (2019) e126207.

[41] T.C.S. Ferreira, I.P. Sauter, L. Borda-Samper, E. Bentivoglio, J.P. DaMata, N.N. Taniwaki, P.R. Orrego, J.E. Araya, N. Lincopan, M. Cortez, Effect of DODAB Nano-Sized Cationic Bilayer Fragments against Leishmania amazonensis, Molecules (Basel, Switzerland) 25(23) (2020) 5741.

[42] J.S.P. Doehl, J. Sádlová, H. Aslan, K. Pružinová, S. Metangmo, J. Votýpka, S. Kamhawi, P. Volf, D.F. Smith, Leishmania HASP and SHERP Genes Are Required for In Vivo Differentiation, Parasite Transmission and Virulence Attenuation in the Host, PLoS Pathog 13(1) (2017) e1006130-e1006130.

[43] J.A. Vizcaíno, A. Csordas, N. del-Toro, J.A. Dianes, J. Griss, I. Lavidas, G. Mayer, Y. Perez-Riverol, F. Reisinger, T. Ternent, Q.W. Xu, R. Wang, H. Hermjakob, 2016 update of the PRIDE database and its related tools, Nucleic acids research 44(D1) (2016) D447-56.

[44] M.J. Alves, R. Kawahara, R. Viner, W. Colli, E.C. Mattos, M. Thaysen-Andersen, M.R. Larsen, G. Palmisano, Comprehensive glycoprofiling of the epimastigote and trypomastigote stages of Trypanosoma cruzi, Journal of proteomics 151 (2017) 182–192.

[45] G.S. de Oliveira, R. Kawahara, L. Rosa-Fernandes, S.N. Mule, C.C. Avila, M.M.G. Teixeira, M.R. Larsen, G. Palmisano, Development of a Trypanosoma cruzi strain typing assay using MS2 peptide spectral libraries (Tc-STAMS2), PLoS neglected tropical diseases 12(4) (2018) e0006351.

[46] G.S. de Oliveira, R. Kawahara, L. Rosa-Fernandes, C.C. Avila, M.R. Larsen, J.M. Pereira Alves, G. Palmisano, Novel DNA coding regions and protein arginylation reveal unexplored T. cruzi proteome and PTMs, International Journal of Mass Spectrometry 418 (2017) 51–66.

[47] Tyanova S, Temu T, Sinitcyn P, Carlson A, Hein MY, Geiger T, Mann M, C. J., The Perseus computational platform for comprehensive analysis of (prote)omics data, Nature methods 13(9) (2016) 731–40.

[48] K.J. Livak, T.D. Schmittgen, Analysis of relative gene expression data using real-time quantitative PCR and the 2(-Delta Delta C(T)) Method, Methods (San Diego, Calif.) 25(4) (2001) 402–8.

[49] Z. Ren, J. Chen, R.A. Khalil, Zymography as a Research Tool in the Study of Matrix Metalloproteinase Inhibitors, Methods in molecular biology (Clifton, N.J.) 1626 (2017) 79–102.

[50] P. Salotra, R. Ralhan, G. Sreenivas, Heat-stress induced modulation of protein phosphorylation in virulent promastigotes of Leishmania donovani, Int J Biochem Cell Biol 32(3) (2000) 309–16.

[51] D. Rosenzweig, D. Smith, P.J. Myler, R.W. Olafson, D. Zilberstein, Post-translational modification of cellular proteins during Leishmania donovani differentiation, Proteomics 8(9) (2008) 1843–50.

[52] M.A. Morales, R. Watanabe, M. Dacher, P. Chafey, J. Osorio y Fortéa, D.A. Scott, S.M. Beverley, G. Ommen, J. Clos, S. Hem, P. Lenormand, J.C. Rousselle, A. Namane, G.F. Späth, Phosphoproteome dynamics reveal heat-shock protein complexes specific to the Leishmania donovani infectious stage, Proceedings of the National Academy of Sciences of the United States of America 107(18) (2010) 8381–6.

[53] G.F. Späth, S. Drini, N. Rachidi, A touch of Zen: post-translational regulation of the Leishmania stress response, Cell Microbiol 17(5) (2015) 632–8.

[54] P. Kaur, M. Garg, A. Hombach-Barrigah, J. Clos, N. Goyal, MAPK1 of Leishmania donovani interacts and phosphorylates HSP70 and HSP90 subunits of foldosome complex, Sci Rep 7(1) (2017) 10202.

[55] S. Sundar, B. Singh, Understanding Leishmania parasites through proteomics and implications for the clinic, Expert Rev Proteomics 15(5) (2018) 371–390.

[56] Ecological and evolutionary pressures on leishmanial parasites, Brazilian Journal of Genetics 20 (1997).

[57] N.G. Saravia, I. Segura, A.F. Holguin, C. Santrich, L. Valderrama, C. Ocampo, Epidemiologic, genetic, and clinical associations among phenotypically distinct populations of Leishmania (Viannia) in Colombia, The American journal of tropical medicine and hygiene 59(1) (1998) 86–94.

[58] E. Cupolillo, L.R. Brahim, C.B. Toaldo, M.P. de Oliveira-Neto, M.E. de Brito, A. Falqueto, M. de Farias Naiff, G. Grimaldi, Jr., Genetic polymorphism and molecular epidemiology of Leishmania (Viannia) braziliensis from different hosts and geographic areas in Brazil, Journal of clinical microbiology 41(7) (2003) 3126–32.

[59] K.P. Chang, B.S. McGwire, Molecular determinants and regulation of Leishmania virulence, Kinetoplastid biology and disease 1(1) (2002) 1.

[60] M. Rossi, N. Fasel, How to master the host immune system? Leishmania parasites have the solutions!, International immunology 30(3) (2018) 103–111.

[61] M. Cortez, C. Huynh, M.C. Fernandes, K.A. Kennedy, A. Aderem, N.W. Andrews, Leishmania promotes its own virulence by inducing expression of the host immune inhibitory ligand CD200, Cell host & microbe 9(6) (2011) 463–71.

[62] T.C.S. Ferreira, I.P. Sauter, L. Borda-Samper, E. Bentivoglio, J.P. DaMata, N.N. Taniwaki, P.R. Orrego, J.E. Araya, N. Lincopan, M. Cortez, Effect of DODAB Nano-Sized Cationic Bilayer Fragments against Leishmania amazonensis, Molecules (Basel, Switzerland) 25(23) (2020).

[63] F. Real, M. Pouchelet, M. Rabinovitch, Leishmania (L.) amazonensis: fusion between parasitophorous vacuoles in infected bone-marrow derived mouse macrophages, Experimental parasitology 119(1) (2008) 15–23.

[64] F. Real, R.A. Mortara, The diverse and dynamic nature of Leishmania parasitophorous vacuoles studied by multidimensional imaging, PLoS neglected tropical diseases 6(2) (2012) e1518.

[65] N.G. Saravia, M.A. Gemmell, S.L. Nance, N.L. Anderson, Two-dimensional electrophoresis used to differentiate the causal agents of American tegumentary leishmaniasis, Clinical chemistry 30(12 Pt 1) (1984) 2048-52.

[66] H. Hajjaran, M. Mohammadi Bazargani, M. Mohebali, R. Burchmore, G.H. Salekdeh, E. Kazemi-Rad, M.R. Khoramizadeh, Comparison of the Proteome Profiling of Iranian isolates of Leishmania tropica, L. major and L. infantum by Two-Dimensional Electrophoresis (2-DE) and Mass-spectrometry, Iranian journal of parasitology 10(4) (2015) 530-40.

[67] M.A. Lynn, A.K. Marr, W.R. McMaster, Differential quantitative proteomic profiling of Leishmania infantum and Leishmania mexicana density gradient separated membranous fractions, Journal of proteomics 82 (2013) 179–92.

[68] N. Acestor, S. Masina, J. Walker, N.G. Saravia, N. Fasel, M. Quadroni, Establishing two-dimensional gels for the analysis of Leishmania proteomes, Proteomics 2(7) (2002) 877–9.

[69] M. Bente, S. Harder, M. Wiesgigl, J. Heukeshoven, C. Gelhaus, E. Krause, J. Clos, I. Bruchhaus, Developmentally induced changes of the proteome in the protozoan parasite Leishmania donovani, Proteomics 3(9) (2003) 1811–29.

[70] J.I. Aoki, S.M. Muxel, R.A. Zampieri, M.F. Laranjeira-Silva, K.E. Müller, A.H. Nerland, L.M. Floeter-Winter, RNA-seq transcriptional profiling of Leishmania amazonensis reveals an arginase-dependent gene expression regulation, PLoS neglected tropical diseases 11(10) (2017) e0006026.

[71] D. Rosenzweig, D. Smith, F. Opperdoes, S. Stern, R.W. Olafson, D. Zilberstein, Retooling Leishmania metabolism: from sand fly gut to human macrophage, FASEB journal: official publication of the Federation of American Societies for Experimental Biology 22(2) (2008) 590–602.

[72] J. Drummelsmith, V. Brochu, I. Girard, N. Messier, M. Ouellette, Proteome mapping of the protozoan parasite Leishmania and application to the study of drug targets and resistance mechanisms, Molecular & cellular proteomics: MCP 2(3) (2003) 146–55.

[73] S.K. Gupta, B.S. Sisodia, S. Sinha, K. Hajela, S. Naik, A.K. Shasany, A. Dube, Proteomic approach for identification and characterization of novel immunostimulatory proteins from soluble antigens of Leishmania donovani promastigotes, Proteomics 7(5) (2007) 816–23.

[74] A. Kumar, P. Misra, B. Sisodia, A.K. Shasany, S. Sundar, A. Dube, Proteomic analyses of membrane enriched proteins of Leishmania donovani Indian clinical isolate by mass spectrometry, Parasitology international 64(4) (2015) 36–42.

[75] P. Tsigankov, P.F. Gherardini, M. Helmer-Citterich, G.F. Späth, D. Zilberstein, Phosphoproteomic analysis of differentiating Leishmania parasites reveals a unique stage-specific phosphorylation motif, Journal of proteome research 12(7) (2013) 3405–12.

[76] P.J. Alcolea, A. Alonso, V. Larraga, Proteome profiling of Leishmania infantum promastigotes, The Journal of eukaryotic microbiology 58(4) (2011) 352–8.

[77] D.M. Forrest, M. Batista, F.K. Marchini, A.J. Tempone, Y.M. Traub-Csekö, Proteomic analysis of exosomes derived from procyclic and metacyclic-like cultured Leishmania infantum chagasi, Journal of proteomics 227 (2020) 103902.

[78] N. Pinho, J.R. Wiśniewski, G. Dias-Lopes, L. Sabóia-Vahia, A.C.S. Bombaça, C. Mesquita-Rodrigues, R. Menna-Barreto, E. Cupolillo, J.B. de Jesus, G. Padrón, P. Cuervo, In-depth quantitative proteomics uncovers specie-specific metabolic programs in Leishmania (Viannia) species, PLoS neglected tropical diseases 14(8) (2020) e0008509.

[79] A.M. Silva, A. Cordeiro-da-Silva, G.H. Coombs, Metabolic variation during development in culture of Leishmania donovani promastigotes, PLoS neglected tropical diseases 5(12) (2011) e1451.

[80] M. Arjmand, A. Madrakian, G. Khalili, A. Najafi Dastnaee, Z. Zamani, Z. Akbari, Metabolomics-Based Study of Logarithmic and Stationary Phases of Promastigotes in Leishmania major by 1H NMR Spectroscopy, Iranian biomedical journal 20(2) (2016) 77–83.

[81] A.E. Ramer-Tait, S.M. Lei, B.H. Bellaire, J.K. Beetham, Differential surface deposition of complement proteins on logarithmic and stationary phase Leishmania chagasi promastigotes, The Journal of parasitology 98(6) (2012) 1109–16.

[82] C. Folgueira, J.M. Requena, A postgenomic view of the heat shock proteins in kinetoplastids, FEMS microbiology reviews 31(4) (2007) 359–77.

[83] T. Sherwin, K. Gull, The cell division cycle of Trypanosoma brucei brucei: timing of event markers and cytoskeletal modulations, Philosophical transactions of the Royal Society of London. Series B, Biological sciences 323(1218) (1989) 573-88.

[84] M.W. Rochlin, M.E. Dailey, P.C. Bridgman, Polymerizing microtubules activate site-directed F-actin assembly in nerve growth cones, Molecular biology of the cell 10(7) (1999) 2309–27.

[85] C.A. Ramírez, J.M. Requena, C.J. Puerta, Alpha tubulin genes from Leishmania braziliensis: genomic organization, gene structure and insights on their expression, BMC genomics 14 (2013) 454.

[86] A.C. Ivens, C.S. Peacock, E.A. Worthey, L. Murphy, G. Aggarwal, M. Berriman, E. Sisk, M.A. Rajandream, E. Adlem, R. Aert, A. Anupama, Z. Apostolou, P. Attipoe, N. Bason, C. Bauser, A. Beck, S.M. Beverley, G. Bianchettin, K. Borzym, G. Bothe, C.V. Bruschi, M. Collins, E. Cadag, L. Ciarloni, C. Clayton, R.M. Coulson, A. Cronin, A.K. Cruz, R.M. Davies, J. De Gaudenzi, D.E. Dobson, A. Duesterhoeft, G. Fazelina, N. Fosker, A.C. Frasch, A. Fraser, M. Fuchs, C. Gabel, A. Goble, A. Goffeau, D. Harris, C. Hertz-Fowler, H. Hilbert, D. Horn, Y. Huang, S. Klages, A. Knights, M. Kube, N. Larke, L. Litvin, A. Lord, T. Louie, M. Marra, D. Masuy, K. Matthews, S. Michaeli, J.C. Mottram, S. Müller-Auer, H. Munden, S. Nelson, H. Norbertczak, K. Oliver, S. O’Neil, M. Pentony, T.M. Pohl, C. Price, B. Purnelle, M.A. Quail, E. Rabbinowitsch, R. Reinhardt, M. Rieger, J. Rinta, J. Robben, L. Robertson, J.C. Ruiz, S. Rutter, D. Saunders, M. Schäfer, J. Schein, D.C. Schwartz, K. Seeger, A. Seyler, S. Sharp, H. Shin, D. Sivam, R. Squares, S. Squares, V. Tosato, C. Vogt, G. Volckaert, R. Wambutt, T. Warren, H. Wedler, J. Woodward, S. Zhou, W. Zimmermann, D.F. Smith, J.M. Blackwell, K.D. Stuart, B. Barrell, P.J. Myler, The genome of the kinetoplastid parasite, Leishmania major, Science (New York, N.Y.) 309(5733) (2005) 436-42.

[87] S.M. Landfear, D.F. Wirth, Control of tubulin gene expression in the parasitic protozoan Leishmania enriettii, Nature 309(5970) (1984) 716-7.

[88] K. Leifso, G. Cohen-Freue, N. Dogra, A. Murray, W.R. McMaster, Genomic and proteomic expression analysis of Leishmania promastigote and amastigote life stages: the Leishmania genome is constitutively expressed, Molecular and biochemical parasitology 152(1) (2007) 35–46.

[89] D. Fong, K.P. Chang, Tubulin biosynthesis in the developmental cycle of a parasitic protozoan, Leishmania mexicana: changes during differentiation of motile and nonmotile stages, Proceedings of the National Academy of Sciences of the United States of America 78(12) (1981) 7624–8.

[90] D. Fong, M. Wallach, J. Keithly, P.W. Melera, K.P. Chang, Differential expression of mRNAs for alpha- and beta-tubulin during differentiation of the parasitic protozoan Leishmania mexicana, Proceedings of the National Academy of Sciences of the United States of America 81(18) (1984) 5782–6.

[91] D. Fong, K.P. Chang, Changes in tubulin mRNAs during differentiation of a parasitic protozoan Leishmania mexicana, Annals of the New York Academy of Sciences 466 (1986) 129–31.

[92] M.J. North, G.H. Coombs, Proteinases of Leishmania mexicana amastigotes and promastigotes: analysis by gel electrophoresis, Molecular and biochemical parasitology 3(5) (1981) 293–300.

[93] G.H. Coombs, Proteinases of Leishmania mexicana and other flagellate protozoa, Parasitology 84(1) (1982) 149–55.

[94] M. Klemba, D.E. Goldberg, Biological roles of proteases in parasitic protozoa, Annual review of biochemistry 71 (2002) 275–305.

[95] M. Silva-Almeida, F. Souza-Silva, B.A. Pereira, M.L. Ribeiro-Guimarães, C.R. Alves, Overview of the organization of protease genes in the genome of Leishmania spp, Parasites & vectors 7 (2014) 387.

[96] C.R. Alves, R.S. Souza, K.D.S. Charret, L.M.C. Côrtes, M.P. Sá-Silva, L. Barral-Veloso, L.F.G. Oliveira, F.S. da Silva, Understanding serine proteases implications on Leishmania spp lifecycle, Experimental parasitology 184 (2018) 67–81.

[97] R. Etges, J. Bouvier, C. Bordier, The major surface protein of Leishmania promastigotes is a protease, The Journal of biological chemistry 261(20) (1986) 9098–101.

[98] C. Yao, J.E. Donelson, M.E. Wilson, The major surface protease (MSP or GP63) of Leishmania sp. Biosynthesis, regulation of expression, and function, Molecular and biochemical parasitology 132(1) (2003) 1–16.

[99] M. Hallé, M.A. Gomez, M. Stuible, H. Shimizu, W.R. McMaster, M. Olivier, M.L. Tremblay, The Leishmania surface protease GP63 cleaves multiple intracellular proteins and actively participates in p38 mitogen-activated protein kinase inactivation, The Journal of biological chemistry 284(11) (2009) 6893–908.

[100] C.S. Chang, T.J. Inserra, J.A. Kink, D. Fong, K.P. Chang, Expression and size heterogeneity of a 63 kilodalton membrane glycoprotein during growth and transformation of Leishmania mexicana amazonensis, Molecular and biochemical parasitology 18(2) (1986) 197–210.

[101] C. Bordier, The promastigote surface protease of Leishmania, Parasitology today (Personal ed.) 3(5) (1987) 151–3.

[102] L.L. Button, D.G. Russell, H.L. Klein, E. Medina-Acosta, R.E. Karess, W.R. McMaster, Genes encoding the major surface glycoprotein in Leishmania are tandemly linked at a single chromosomal locus and are constitutively transcribed, Molecular and biochemical parasitology 32(2-3) (1989) 271–83.

[103] L.L. Button, W.R. McMaster, Molecular cloning of the major surface antigen of leishmania, The Journal of experimental medicine 167(2) (1988) 724–9.

[104] L.S. Medina, B.A. Souza, A. Queiroz, L.H. Guimarães, P.R. Lima Machado, M.C. E, M.E. Wilson, A. Schriefer, The gp63 Gene Cluster Is Highly Polymorphic in Natural Leishmania (Viannia) braziliensis Populations, but Functional Sites Are Conserved, PloS one 11(9) (2016) e0163284.

[105] M.M. Guay-Vincent, C. Matte, A.M. Berthiaume, M. Olivier, M. Jaramillo, A. Descoteaux, Revisiting Leishmania GP63 host cell targets reveals a limited spectrum of substrates, PLoS Pathog 18(10) (2022) e1010640.

[106] P. Cuervo, L. Sabóia-Vahia, F. Costa Silva-Filho, O. Fernandes, E. Cupolillo, D.E.J. JB, A zymographic study of metalloprotease activities in extracts and extracellular secretions of Leishmania (Viannia) braziliensis strains, Parasitology 132(Pt 2) (2006) 177–85.

[107] J.M. Requena, A.M. Montalvo, J. Fraga, Molecular Chaperones of Leishmania: Central Players in Many Stress-Related and -Unrelated Physiological Processes, BioMed research international 2015 (2015) 301326.

[108] M.E. Feder, G.E. Hofmann, Heat-shock proteins, molecular chaperones, and the stress response: evolutionary and ecological physiology, Annual review of physiology 61 (1999) 243–82.

[109] J.L. Johnson, Evolution and function of diverse Hsp90 homologs and cochaperone proteins, Biochimica et biophysica acta 1823(3) (2012) 607–13.

[110] A. Rastrojo, L. Corvo, R. Lombraña, J.C. Solana, B. Aguado, J.M. Requena, Analysis by RNA-seq of transcriptomic changes elicited by heat shock in Leishmania major, Scientific Reports 9(1) (2019) 6919.

